# Integrating temporal single-cell gene expression modalities for trajectory inference and disease prediction

**DOI:** 10.1101/2022.03.01.482381

**Authors:** Jolene S. Ranek, Natalie Stanley, Jeremy E. Purvis

**Author notes:** Correspondence to Natalie Stanley, and Jeremy Purvis.

## Abstract

Current methods for analyzing single-cell datasets have relied primarily on static gene expression measurements to characterize the molecular state of individual cells. However, capturing temporal changes in cell state is crucial for the interpretation of dynamic phenotypes such as the cell cycle, development, or disease progression. RNA velocity infers the direction and speed of transcriptional changes in individual cells, yet it is unclear how these temporal gene expression modalities may be leveraged for predictive modeling of cellular dynamics. Here, we present the first task-oriented benchmarking study that investigates integration of temporal sequencing modalities for dynamic cell state prediction. We benchmark eight integration approaches on eight datasets spanning different biological contexts, sequencing technologies, and species. We find that integrated data more accurately infers biological trajectories and achieves increased performance on classifying cells according to perturbation and disease states. Furthermore, we show that simple concatenation of spliced and unspliced molecules performs consistently well on classification tasks and can be used over more memory intensive and computationally expensive methods. This work provides users with practical recommendations for task-specific integration of single-cell gene expression modalities.

## Introduction

Single-cell RNA sequencing (scRNA-seq) technologies have enabled the functional characterization of cellular states associated with dynamic biological processes such as development [1, 2, 3] and disease progression [4, 5, 6, 7]. While transcriptomic information holds great promise for gaining insight into the biological mechanisms that govern phenotypic changes, inference has been traditionally limited to incompletely-sampled static mature mRNA measurements. This poses two fundamental challenges for robust prediction of the dynamic progression of cell state. First, many gene regulatory mechanisms can give rise to the same distribution of mature mRNA measurements [8]. Second, snapshot data often fails to fully capture the large biological variability required for population-level inference by missing important transition states or rare cell populations [9, 10, 11].

More recently, computational tools such as RNA velocity have been used to extract directed dynamic information from single cells [12, 13, 14, 15, 16]. By leveraging unspliced pre mRNA and spliced mature mRNA molecules in a kinetic model, RNA velocity can predict the future transcriptional state of a cell. Indeed, RNA velocity has been successfully incorporated into algorithms for inferring fate probabilities [17], gene regulatory networks [18], differentiation trajectories [19, 20, 21], and embeddings [22]. However, it is still unclear whether integrating spliced gene expression with either unspliced molecules or RNA velocity predictions is useful for predictive modeling at the data-level. Such an integrated approach may help uncover salient features predictive of cell type or response to therapy, enhance our understanding of the relationship between cell states, or provide insight into the molecular pathways that drive a cell’s transition to a more pathological phenotype.

Single-cell multi-omics data integration methods have had great success in fusing different molecular data types, or modalities for disease subtyping, predicting biomarkers, or uncovering cross-modality correlations [23, 24]. Here, integration methods aim to merge individual layers of single-cell data (e.g. transcriptomic, proteomic, epigenomic) into a unified consensus representation, such as an integrated graph [25] or a joint-embedding [24, 26]. To do so, computational approaches have leveraged techniques, including kernel learning [27, 28], matrix factorization [29, 30, 31, 32, 33], or deep learning [34]. Moreover, downstream analysis of integrated multi-omics data has has provided fundamental insights into the molecular mechanisms underlying complex biological processes, including disease heterogeneity and pathological development [35].

Motivated by identifying a new more biologically-meaningful set of features underlying cellular dynamics, we investigate integration of gene expression modalities at three distinct temporal stages of gene regulation: unspliced, spliced, and RNA velocity. We benchmark eight integration approaches on eight biological datasets with applications ranging from cellular differentiation to disease progression. We show that unspliced and spliced integration improves predictive performance when inferring biological trajectories, perturbation conditions, and disease states. This work illustrates how integrated temporal gene expression modalities may be leveraged for predictive modeling of cellular dynamics.

## Results

We compared eight integration approaches for recovering discrete and continuous variation in cell and disease states. In the sections that follow, we will describe the integration results in more detail. We will begin by giving an introduction of the datasets used in this study. Next, we will provide details about the benchmarking design, including the integration methods considered and the evaluation criteria for each prediction task. We will then demonstrate how an integrative analysis can be used to obtain increased biological insight over spliced expression alone. Ultimately, we will end with practical recommendations for task-specific integration.

### Description of datasets

We tested integration method performance on inferring biological trajectories or classifying cells according to perturbation condition or disease status across eight publicly available single-cell RNA sequencing datasets (see Methods, Supplementary Table 1). Datasets were grouped into three general categories according to the prediction task. Here, we briefly introduce the datasets used in this study.

#### Datasets for Trajectory Inference (TI)

We evaluated inference of biological trajectories using two single-cell RNA sequencing datasets representing the cell cycle and stem cell differentiation. To assess inference of cell cycle, we considered a mouse embryonic stem cell cycle dataset [36], where embryonic stem cells were collected along three stages of the cell cycle (G1, S, G2/M). Cell cycle phase was manually annotated *a priori* based on flow sorting cells according to the Hoeschst 33342 stained distribution. The authors of the original study used this dataset to assess the proportion of cell-to-cell heterogeneity that arises from cell cycle variation. To assess inference of a complex differentiation trajectory, we considered a mouse hematopoietic stem and progenitor cell differentiation (HPSC) dataset [37]. Here, the transcriptomes of HPSCs were profiled and nine cell surface protein measurements (Supplementary Table 3) were used to annotate six subpopulations, including, long-term hematopoeitic stem cells (LT-HSC), lymphoid multipotent progenitors (LMPP), multipotent progenitors (MPP), megakaryocyte-erythrocyte progenitors (MEP), common myeloid progenitors (CMP), and granulocyte-monocyte progenitors (GMP). Moreover, in the original study, reconstruction of the differentiation trajectory revealed dynamic gene expression patterns consistent with early lymphoid, erythroid, and granulocyte-macrophage differentiation. For our analysis, cells were excluded if they did not have ground truth annotations.

#### Datasets for perturbation classification

To assess integration performance on classifying cells according to perturbation condition, we considered three diverse datasets with clinical relevance representing drug stimulation and treatment of cells, denoted as LPS stimulation, INF*γ* stimulation, and AML chemotherapy, respectively. In the LPS stimulation dataset [38], RAW 264.7 macrophage-like cells were treated with time course of lipopolysaccharide (0 min, 50 min-, 150min-, 300min-LPS) to induce NF-*κ*B expression. NF-*κ*B is a transcription factor that serves as a master regulator of inflammatory responses from macrophages and other innate immune cells [39]. The authors of this study integrated live cell imaging with single-cell RNA sequencing to demonstrate that NF-*κ*B signaling shapes gene expression and has a functional role on cellular phenotypes. Therefore, in our experiments, we sought to classify cells according to stimulation condition (e.g. 150min-LPS). In the INF*γ* stimulation dataset [40], pancreatic islet cells from three donors were stimulated with or without Interferon-*γ* (INF*γ*) for 24 hours. INF*γ* is a proinflammatory cytokine that has been implicated in pancreatic beta cell damage during the pathogenesis of Type I Diabetes [41]. Here, the authors applied their method MELD to characterize INF*γ* treatment response across pancreatic islet cell populations and identified a non-responsive subpopulation of beta cells characterized by high expression of insulin. Consequently, we sought to classify INF*γ*-stimulated from unstimulated cells.

Lastly, the AML chemotherapy dataset [5] consisted of peripheral blood mononuclear cells (PBMCs) collected from a patient with Acute Myeloid Leukemia (AML) at baseline or after two or four days of treatment with chemotherapy agents Venetoclax and Azacitidine. It is hypothesized that the persistence of leukemia stem cells (LSCs) following treatment drives disease severity, relapse, and results in worse clinical outcomes [7, 42]. Here, the authors demonstrate how chemotherapy treatment induces the depletion of LSCs through metabolic reprogramming, where oxidative phosphorylation, a critical pathway for LSC maintenance and survival, is suppressed. Thus, we sought to classify PBMCs according to treatment condition (day 0, day 2, day 4).

#### Datasets for disease status classification

To assess integration performance on classifying cells according to disease status, we considered three case/ control datasets of two disease systems, Acute Myeloid Leukemia (AML) and Multiple Sclerosis (MS). In the first dataset [7], Leukemia stem cells (LSCs) were collected from AML patients at treatment-naive diagnosis (*N* = 5) and following relapse after chemotherapy treatment (*N* = 5). Here, the authors compared diagnosis from relapse samples to characterize gene expression heterogeneity during AML disease progression and show that differences were largely due to metabolic reprogramming, apoptotic signaling, and chemokine signaling. Therefore, in our experiments, we sought to classify diagnosis from relapse cells. For the second and third study, we considered a Multiple Sclerosis dataset [6], where PBMCs and cerebral spinal fluid (CSF) were collected from MS patients (*N* = 5) and controls (*N* = 5). MS is a chronic inflammatory disorder of the central nervous system that results in neurological dysfunction [43]. When examining the transcriptional profiles of MS patient cells as compared to controls, CSF exhibited differences in cell type composition, including an enrichment of myeloid dendritic cells and the expansion of CD4+ cytotoxic T cells and late stage B cells. In contrast, PBMCs exhibited increased transcriptional diversity with an increased proportion of differentially expressed genes. Consequently, we sought to classify control from MS cells across patients using either CSF or PBMC biological samples.

### Selection of integration methods

The power of multi-omics data integration methods lies in their ability to combine individual layers of data (e.g. spliced expression, RNA velocity) to identify a new set of cellular features that more holistically represents cell type or functional state [23, 44]. Once computed, these features can be used in machine learning models to jointly analyze cell type-specific differences or to obtain clinically meaningful predictions that can inform therapeutics [45, 46]. In this study, our goal is to compare integration approaches for merging gene expression data matrices across the same set of profiled cells in order to evaluate their performance on downstream analysis tasks, including trajectory inference or sample-associated classification of cells. Given the large variety of different integration approaches, we performed a systematic evaluation of eight integration methods by selecting and grouping approaches according to two categories: early integration approaches and intermediate integration approaches. First, we consider early integration approaches as baseline computational strategies for merging individual modalities into one input matrix. Here, we selected three representative baseline strategies (cell-wise concatentation, cell-wise sum, CellRank [17]), in addition to an unintegrated control. In contrast, we consider intermediate integration approaches as computational strategies that transform individual modalities into an intermediate representation prior to merging, such as a cell similarity graph or a subspace. Within this category, we selected four representative methods, including Similarity Network Fusion (SNF) [25], Grassmann Joint Embedding [26], integrated diffusion [24], and Patient Response Estimation Corrected by Interpolation of Subspace Embeddings (PRECISE) [47]. Here, we briefly define the eight integration approaches evaluated in this study. For more details on the overall problem formulation and integration method implementation, see the integration section in the Methods.

1. *Unintegrated:* A representation consisting of one data modality. In this case, our unintegrated data consists of mature spliced expression counts, as this is what is traditionally used for downstream single-cell analysis, as outlined by current best practices [48].
2. *Concatenation:* Modalities are merged through cell-wise concatenation of data matrices.
3. *Sum:* Modalities are merged through summing data matrices.
4. *CellRank:* CellRank [17] merges data modalities by computing a weighted sum of gene expression similarity and RNA velocity transition matrices. We refer to this approach as an early integration strategy as it simply reweights the edges of the original gene expression cell similarity graph according to RNA velocity transition probabilities. Notably, this method is specific to integrating RNA velocity data.
5. *Similarity Network Fusion (SNF):* SNF [25] merges data by first computing an cell affinity graph for each data type. Next, individual modality networks are merged through nonlinear diffusion iterations to obtain a fused network.
6. *Grassmann Joint Embedding:* Grassmann Joint Embedding [26] integrates data modalities by first computing an cell affinity graph for each data modality, and then merges networks through subspace analysis on a Grassmann manifold.
7. *Integrated diffusion:* Integrated diffusion [24] merges data modalities by first computing a diffusion operator for each denoised data type. Next, individual operators are merged by computing a joint diffusion operator.
8. *Patient Response Estimation Corrected by Interpolation of Subspace Embeddings (PRECISE):* PRECISE merges data by first performing principal components analysis (PCA) on each individual modality. Next, principal components are geometrically aligned and consensus features are determined through interpolation. For this analysis, we implement two versions by projecting spliced expression onto (1) the principal vectors (denoted as PRECISE) or (2) the consensus features (denoted as PRECISE consensus).

### Benchmarking overview

Given that gene expression modalities are collected along a temporal axis of gene regulation, we evaluated the performance of integrating unspliced, spliced, or RNA velocity modalities on predicting discrete and continuous variation in cell and disease states across a range of biological scenarios (Supplementary Table 1). Following transcriptomic profiling, spliced and unspliced counts were preprocessed and jointly batch effect-corrected prior to RNA velocity estimation (see Methods, Supplementary Table 1, Supplementary Figures 1-10). For each set of modalities (spliced and unspliced counts, moments of spliced and RNA velocity), our goal is to identify a consensus representation that we can use as input to a predictive model (Figure 1A). We benchmarked eight integration approaches for combining these gene expression modalities by evaluating how well integrated features infer biological trajectories, classify a cell’s response to a drug perturbation, or classify the disease status of a cell. Moreover, to quantify the predictive performance of an integration strategy, we computed several metrics for each prediction task. To assess the quality of trajectory inference prediction following integration, we computed a trajectory inference correlation score to a ground truth reference that takes into account cellular positioning and trajectory-specific dynamically expressed genes. To assess classification performance, we computed the accuracy of predicted labels from an integration strategy using three complementary metrics, such as F1 score, balanced accuracy, and area under the receiver operator curve. For integration methods that required user-specified input parameters (Supplementary Table 2), we performed hyperparameter tuning to select the best performers. We then ranked the overall predictive performance of integration strategies for each task by averaging scores across all datasets (see Methods). This measures how well incorporating dynamic mRNA information aids in recovering intermediate transitions or classifying the state of a cell.

**Figure 1:**
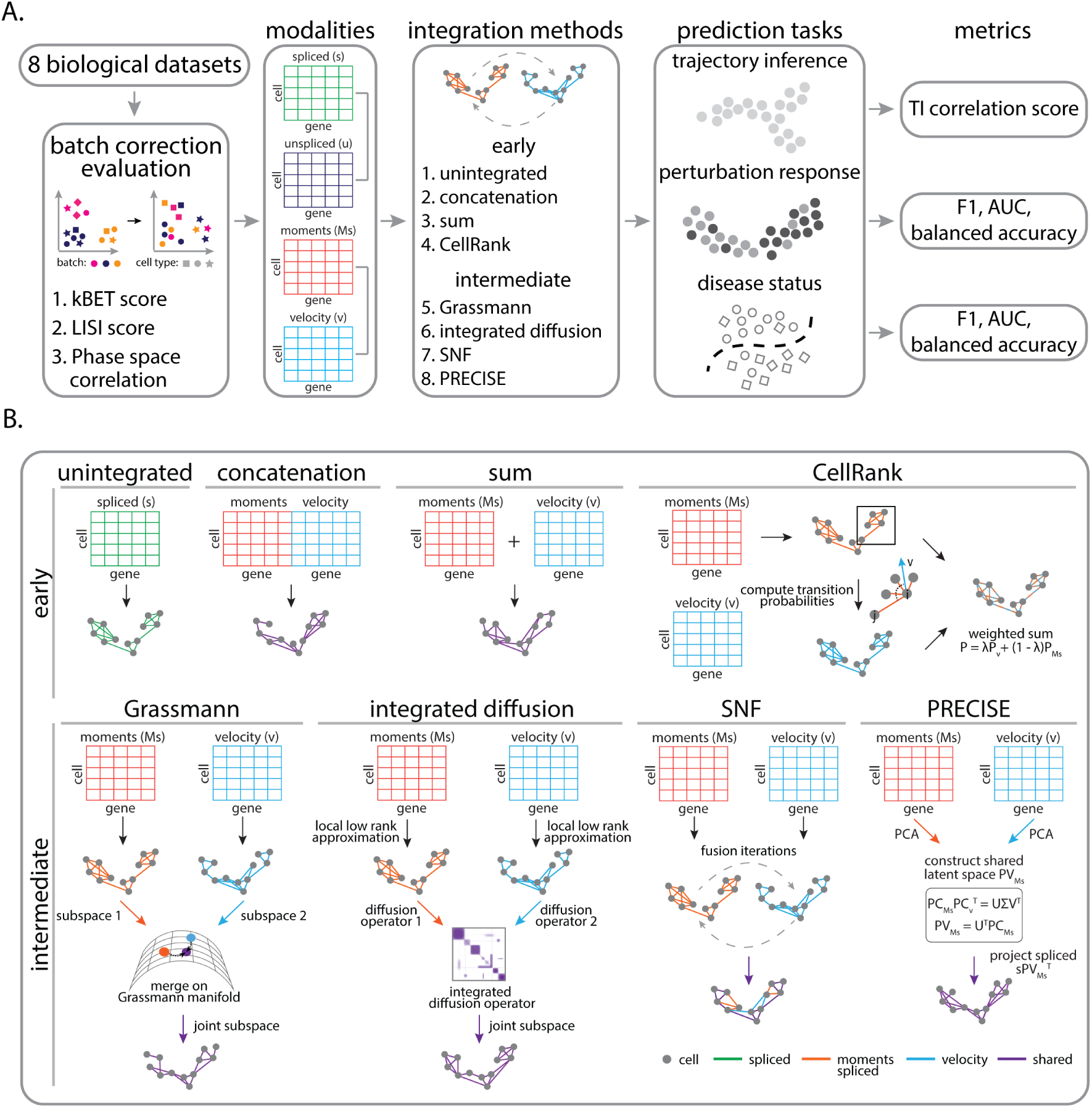
Schematic overview of benchmarking design. (A) Workflow of integration method evaluation. Eight integration approaches and four temporal mRNA modalities are evaluated on three prediction tasks. Data are first preprocessed and jointly batch effect corrected. Next cross-modality integration (spliced and unspliced counts or moments of spliced and RNA velocity) is performed using eight different integration approaches. Features specified through the integration strategy are used to infer trajectories, predict response to drug treatment, and classify patient cells. (B) Overview of data integration strategies (unintegrated, concatenation, sum, CellRank [17], Grassmann joint embedding [26], integrated diffusion [24], SNF [25], and PRECISE [47]).

In selecting an appropriate data integration strategy, it is crucial that the approach is able to satisfy computational challenges that are specific to each modality. First, a method must be robust to varying amounts of sparsity between data types. Single-cell RNA sequencing modalities produce matrices which contain a large proportion of zeros, where only a small fraction of total transcripts are detected due to capture inefficiency, amplification noise, and stochasticity [49]. This sparsity is far greater in unspliced molecules due to polyadenylation enrichment in library preparation [12]. Moreover, given that unspliced, spliced, and RNA velocity predictions are influenced by biological and technical noise, a method must be able to resolve noisy signals for robust prediction. To address these challenges, we compared two classes of integration approaches for combining temporal sequencing modalities, including early integration approaches (concatenation, sum, CellRank) and intermediate integration approaches (Grassmann joint embedding, integrated diffusion, SNF, PRECISE) (see Selection of integration methods, Methods, Figure 1B).

### Integration performance on inference of biological trajectories

When undergoing dynamic processes such as differentiation, cells exhibit a continuum of cell states with fate transitions marked by external stimuli, cell-cell interactions, and stochastic gene expression [50]. One limitation of trajectory inference (TI) reconstruction from snapshot single-cell data is the fact that many gene regulatory mechanisms and cellular dynamics could give rise to the same distribution of cell states [8]. We reasoned that incorporation of unspliced counts or RNA velocity data may provide increased granularity of the state space to more accurately recapitulate the underlying trajectory. To test this hypothesis, we evaluated integration method performance on inferring two types of biological trajectories, cell cycle and differentiation, by measuring their ability to (1) recover known cell population transitions and (2) infer lineage-specific dynamically expressed genes.

In order to construct reference trajectories for evaluation, we chose well-studied biological systems and selected datasets that had gold standard cell type annotations according to the expression of particular characteristic phenotypic markers. Therefore, we selected datasets consisting of mouse embryonic stem cell cycle and mouse hematopoietic stem and progenitor cell differentiation trajectories (see Description of datasets, Methods). We then quantified how well integrated features recapitulated cell cycle or differentiation trajectories by adapting an approach previously used to assess the accuracy of trajectory inference methods [51] (see Methods). Briefly, we constructed predicted trajectories for each integration approach by applying partition-based graph abstraction (PAGA) [52], a state-of-the-art trajectory inference method, on the joint graph following integration. First, PAGA was used on the integrated *k*-nearest neighbor graph to determine directed weighted edges between known cell types according to FACS annotations. Here, the edge weights quantify the strength in connectivity between cell populations, which represents the overall confidence of a cell population transition. Next, we applied diffusion pseudotime [53] to determine an individual cell’s progression through those high-confidence paths. Since integrated features are used as input, the inferred trajectory now contains transcriptional information from a transitional process at or following a measured time point. To assess the accuracy of predicted trajectories, we defined a trajectory inference correlation score that compares predicted trajectories to a ground truth reference trajectory we curated from the literature (see Methods). By taking into account a cell’s position along the trajectory, as well as the features that are dynamically expressed, this correlation metric reflects how well integration infers known cellular dynamics. Moreover, to ensure a robust comparison across integration approaches, we generated predicted trajectories and correlation scores with respect to the same ten random root cells selected from the annotated root cluster (mouse embryonic cell cycle: G1, mouse hematopoietic differentiation: long-term hematopoietic stem cells (LT-HSC)).

When comparing predicted trajectories across integration approaches, we found spliced and unspliced as well as moments of spliced and RNA velocity integrated features led to a higher trajectory inference correlation score when compared to unintegrated data (Figure 2A). For cell cycle, the best performing median TI correlation scores were 0.849, 0.856, 0.750 for unspliced integration, velocity integration, and unintegrated data, respectively. For hematopoietic stem cell differentiation, the scores were 0.792, 0.787, 0.579 for unspliced integration, velocity integration, and unintegrated data, respectively. We next investigated how incorporating temporal gene expression modalities alters the inferred PAGA trajectories and diffusion map embeddings for the top integration performers with respect to unintegrated data (Figure 2B). When examining the PAGA graphs, we found that all predicted trajectories captured the major cell state transitions supported by the literature. For mouse embryonic cell cycle, predicted trajectories included the cyclical transition through the proliferative phases of the cell cycle [36]. For mouse hematopoiesis, predicted trajectories inferred known developmental lineages, with cells transitioning from the multipotent progenitor (MPP) population to early lymphoid (LMPP), erythroid (MEP), and granulocyte-macrophage (GMP) cell populations [54, 37]. In addition to capturing known transitions, predicted trajectories with integrated data resulted in an improved recovery of cellular dynamics. For example, integration of spliced and unspliced counts with SNF better resolves the smooth cyclical progression through the embryonic cell cycle, with cells following a clear trajectory from G1 to S to G2/M (Figure 2B). Moreover, by comparing the change in PAGA connectivity across the same integration strategy for different input modalities (Figure 2B), we observe how temporal gene expression modalities influences the confidence of an inferred cell state transition. When integrating unspliced and spliced features for cell cycle inference, we observe an increase in PAGA connectivity from G2/M to G1 to S phases, whereas RNA velocity integration illustrates the next time point and provides stronger transition weights from G1 to S to G2/M. This added layer of granularity demonstrates prioritized cell type transitions with respect to the underlying gene expression dynamics, which may provide additional insight into the gene regulatory programs that drive specific paths of temporal variation. Lastly, by aggregating trajectory inference correlation scores across datasets, we find integrated diffusion and similarity network fusion amongst the best ranking methods for predicting trajectories with both sets of modalities (Supplementary Figure 11). Taken together, these results indicate that integrating gene expression data improves the ability to predict temporal changes in gene expression along progressive changes in cell state.

**Figure 2:**
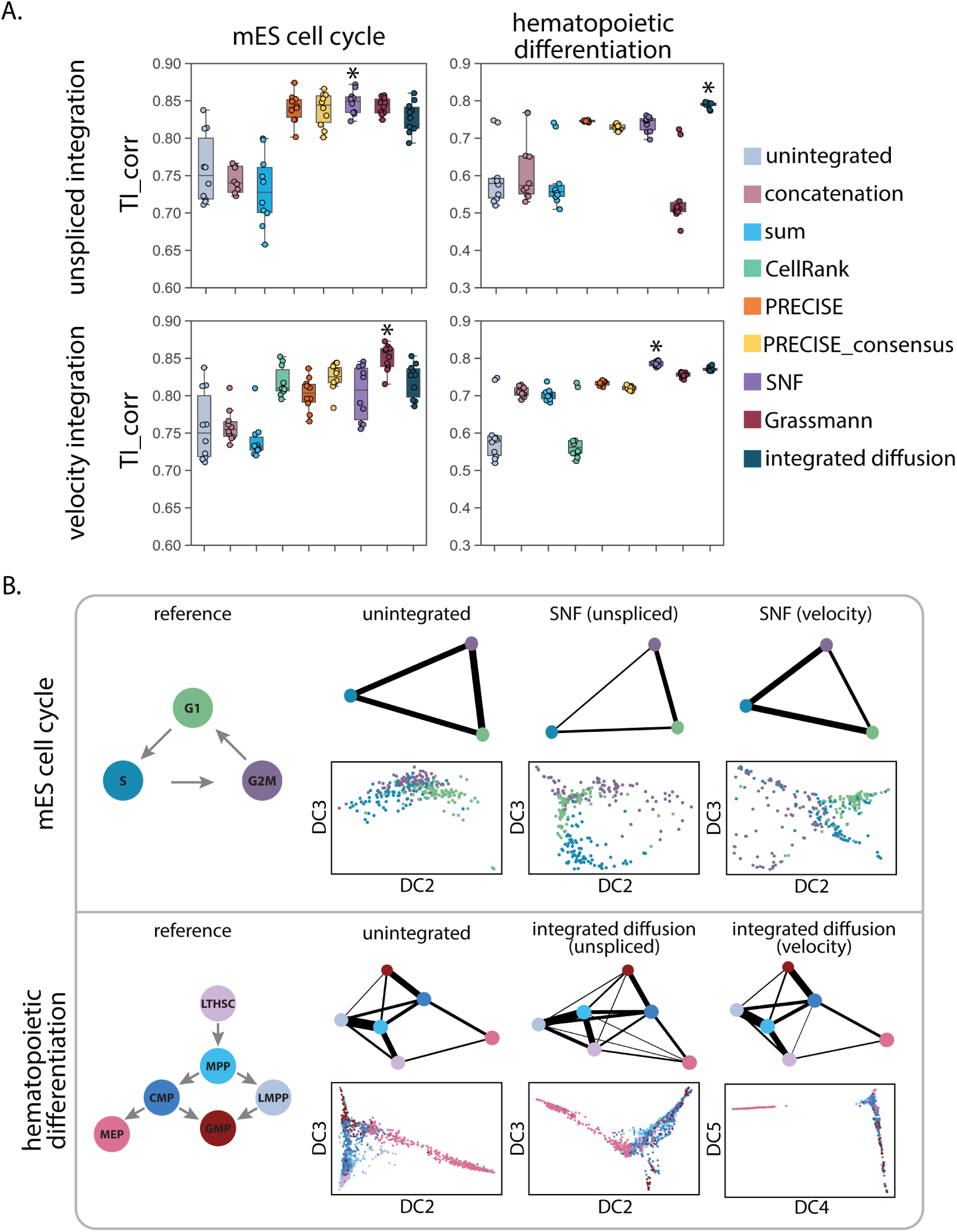
Integration improves inference of cell cycle and differentiation trajectories. Trajectory inference was performed to assess the quality of inferred mouse embryonic cell cycle and mouse hematopoiesis differentiation trajectories from (A top panel) spliced and unspliced or (A bottom panel) moments of spliced and RNA velocity integrated features generated from eight integration methods. The boxplots represent trajectory inference correlation scores (TI_corr_) for ten random root cells. * indicates the method with the highest median TI_corr_ score. (B) PAGA predicted trajectories and diffusion map embeddings representing the inferred biological trajectory for unintegrated data, as well as high ranking performers for unspliced and RNA velocity integration.

### Testing integration under perturbation conditions

A key application of scRNA sequencing is the ability to identify subpopulations of cells that are either responsive or resistant to drug therapy [55]. To examine if integration of unspliced or RNA velocity data can aid in these tasks, we tested integration performance on classifying perturbation condition labels from three diverse datasets with clinical relevance, including lipopolysaccharide (LPS) stimulated macrophage-like cells, Interferon*γ* (INF*γ*) stimulated pancreatic islet cells, and peripheral blood mononuclear cells (PBMCs) collected from a patient with Acute Myeloid Leukemia (AML) after chemotherapy treatment (see Description of datasets). Using these perturbation datasets, we constructed a set of integrated features corresponding to a cell’s transcriptional response following a perturbation. We then considered the problem of cell state classification, where our goal is to learn the annotated condition labels (e.g. INF*γ* stimulated or unstimulated) from the underlying feature set. We labeled or classified cells using label propagation [56] (see Methods) and compared predictions to the ground truth labels using three complementary accuracy metrics, including area under the receiver operator curve (AUC), F1 score, and balanced accuracy (acc_b_). Across all three datasets, we found that integration of spliced and unspliced counts led to higher classification accuracy than unintegrated data (Figure 3A), with median AUCs (best performing integrated: 0.905, 0.953, 0.785; unintegrated: 0.895, 0.930, 0.768) for LPS, INF*γ*, AML chemotherapy datasets, respectively. In contrast, we found that RNA velocity integration generally led to worse classification accuracy than unintegrated data (Figure 3B). One notable exception was integration performed with CellRank, which resulted in a similar performance to unintegrated data, with median AUCs (CellRank: 0.895, 0.934, 0.766, unintegrated: 0.895, 0.930, 0.768). Similar results were obtained for additional metrics, such as F1 score and balanced accuracy (Supplementary Figure 12). As a secondary validation, we trained a support vector machine (SVM) classifier to learn perturbation labels from the shared lower dimensional space following integration. We performed nested 10-fold cross validation to obtain a distribution of predictions for each method and dataset (see Methods). We observed similar classification results with unspliced integration outperforming unintegrated data (Supplementary Figure 13).

**Figure 3:**
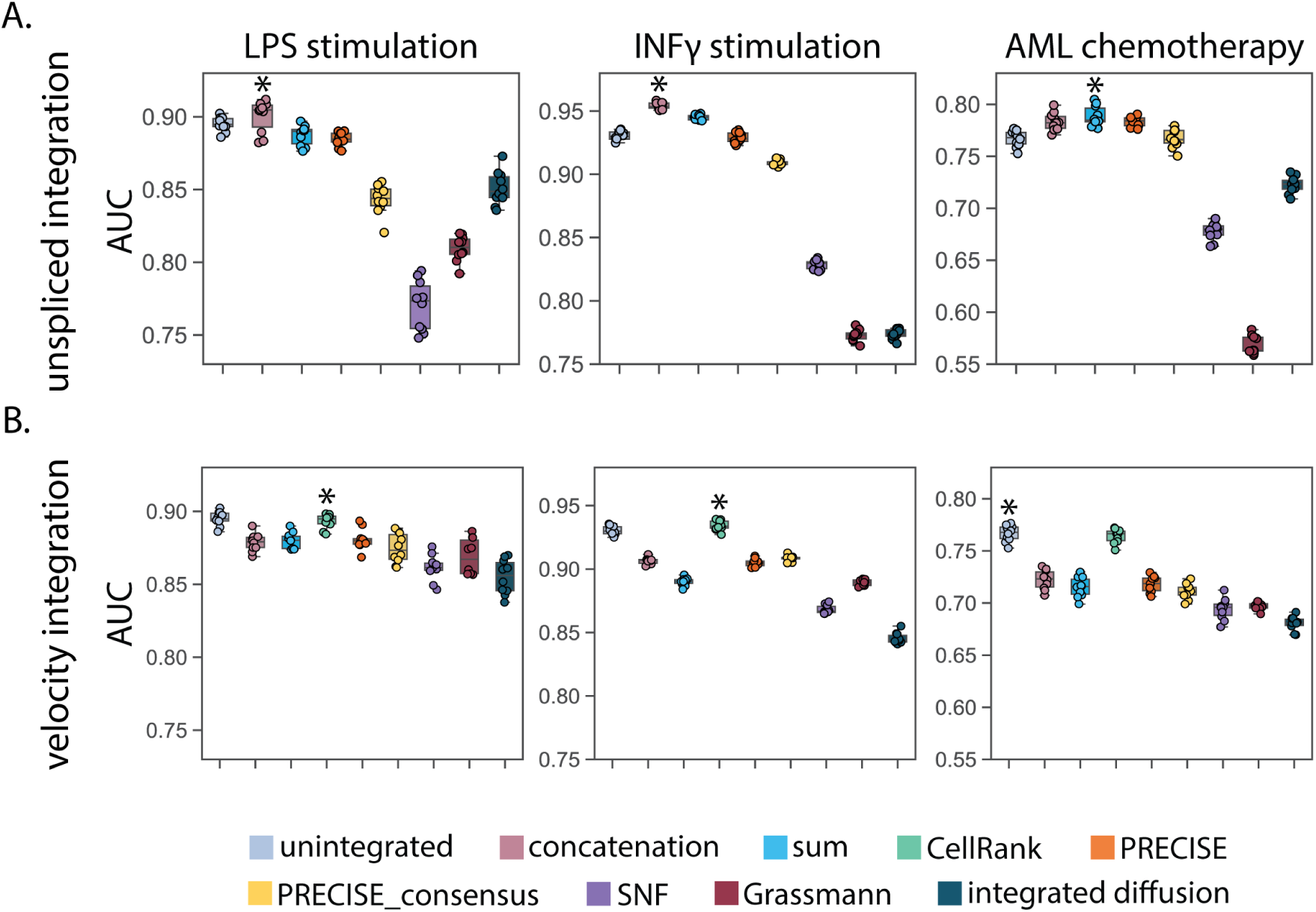
Integrating spliced and unspliced counts improves drug treatment condition prediction. Label propagation was used to classify cells according to treatment response from (A) spliced and unspliced or (B) moments of spliced and RNA velocity integrated features generated from eight integration approaches. The boxplots represent classification accuracy according to area under the receiver operator curve (AUC) and the * represents the method with the highest median AUC. Across all three datasets, spliced and unspliced integration achieves increased classification accuracy over unintegrated data.

To rank methods according to how accurately they can predict a cell’s perturbation, we computed aggregate scores by taking the mean of individual method scores across datasets (see Methods). Overall, we found that early integration strategies (concatenation, sum, CellRank) as well as PRECISE tended to outperform intermediate embedding-based approaches (SNF, Grassmann joint embedding, integrated diffusion) (Supplementary Figure 14). The best performing method for unspliced integration was concatenation (Supplementary Figure 14A), whereas the best performing method for RNA velocity integration was CellRank (Supplementary Figure 14B). Overall, these results suggest that a straightforward integration of spliced and unspliced counts may provide the best strategy to most accurately predict a cell’s associated perturbation. Furthermore, these results illustrate how an integrated analysis of gene expression modalities may provide the granularity necessary for better identifying cells that are strongly associated with a particular treatment condition, which may provide insights into the biological mechanisms conferring a phenotypic response.

### Spliced and unspliced integration improves disease state classification

We next asked if an integrative analysis of unspliced or RNA velocity data can help distinguish discrete disease cell states. In particular, we aimed to evaluate integration performance on predicting whether or not cells were from control or disease patients using three datasets, including an Acute Myeloid Leukemia (AML) diagnosis/relapse dataset, a Multiple Sclerosis (MS) case/control dataset of cerebral spinal fluid (CSF), and a MS case/ control dataset of peripheral blood mononuclear cells (PBMCs) (see Description of datasets). To test whether leveraging temporal gene expression modalities can aid in this tasks, we used the same label propagation strategy; however, now formulated as a binary classification task based on the disease status labels for each cell. Similar to the perturbation results, we found that unspliced integration achieves higher classification accuracy for predicting disease status, with the median AUCs for the best performing methods (0.916, 0.861, 0.884) compared to unintegrated data (0.895, 0.828, 0.825) for AML, MS-CSF, and MS-PBMC datasets, respectively (Figure 4A). Interestingly, we observe differences in the predictive performance of integrated modalities across biological samples (CSF, PBMCs) collected from the same cohort of patients. Overall trends for integration performance were consistent across additional metrics and classifiers (Supplementary Figure 13, Supplementary Figure 15). When ranking each particular method’s performance on classifying the disease status of a cell across datasets, we found the best performing methods for unspliced integration to be PRECISE, sum and concatenation (Supplementary Figure 16).

**Figure 4:**
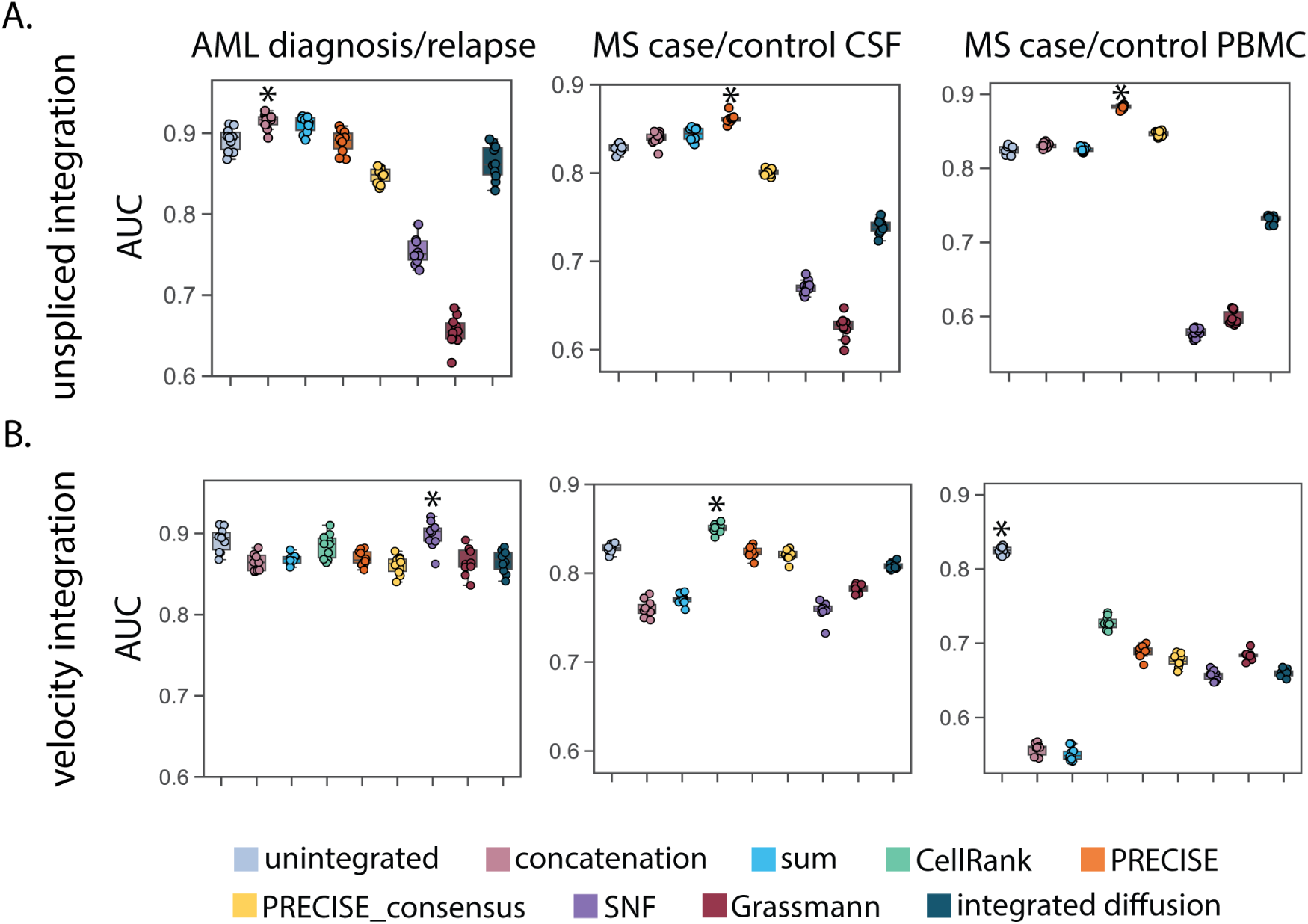
Integrating spliced and unspliced counts improves disease state classification. Label propagation was used to classify cells according to patient disease status from (A) spliced and unspliced or (B) moments of spliced and RNA velocity integrated features generated from eight integration approaches. The boxplots represent classification accuracy according to area under the receiver operator curve (AUC) and the * represents the method with the highest median AUC. Across all three datasets, spliced and unspliced integration achieves increased classification accuracy over unintegrated data.

### Overall integration method performance across datasets and tasks

Figure 5 displays the overall ranked aggregate scores for each method colored according to task (green: trajectory inference, pink: perturbation classification, blue: disease state classification). Across all three tasks, we found unspliced integration (Figure 5A) to be more predictive of cellular state than RNA velocity integration (Figure 5B) or no integration (unintegrated Figure 5A, 5B). While integration method performance varied across datasets, experimental modalities, and tasks, some clear trends emerged. When inferring biological trajectories, unspliced integration with integrated diffusion and similarity network fusion (SNF) provided the highest trajectory inference correlation score to the ground truth (Figure 5A). In comparison, when evaluating perturbation or disease cell state classification, concatenation, sum, and PRECISE were amongst the best ranking methods across all three metrics and datasets (Figure 5A). Collectively, these results indicate that integration method performance is task-specific, with intermediate embedding-based approaches outperforming unintegrated data on inferring biological trajectories and early baseline approaches achieving increased classification performance.

**Figure 5:**
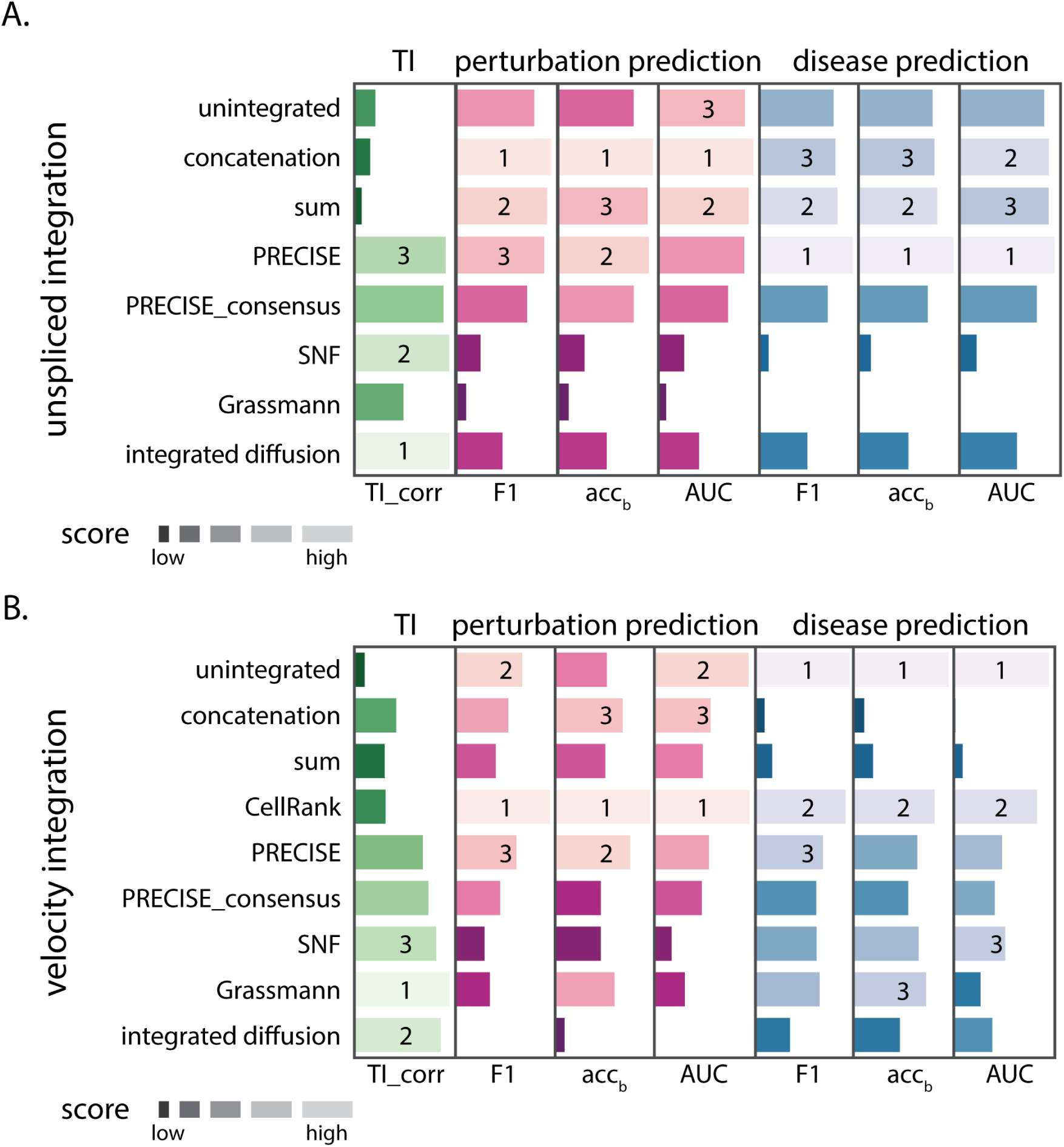
Ranked integration method performance across prediction tasks. Integration methods were ranked by averaging their overall performance across datasets for each prediction task (trajectory inference: green, perturbation classification: blue, and classification of disease status: pink). Ranked scores were computed for several metrics for evaluating a prediction task: (TI_corr_), F1 score, balanced accuracy (acc_b_), and area under the receiver operator curve (AUC)). Here, higher ranked method scores are indicated by a longer lighter bar. (A) Overall quality of spliced and unspliced integration performance according to several metrics for evaluating prediction tasks. (B) Overall quality of moments of spliced and RNA velocity integration performance according to several metrics for evaluating prediction tasks. Of note, CellRank was not performed on unspliced and spliced integration, as it relies on RNA velocity data. Across all three prediction tasks, unspliced integration outperforms unintegrated data, while RNA velocity integration achieves increased trajectory inference correlation and perturbation classification scores.

## Discussion

Here, we investigated integration of unspliced, spliced, and RNA velocity gene expression modalities for resolving discrete and continuous variation in cell and disease states. We found that integrating modalities along a temporal axis of gene regulation provides additional information necessary for robustly predicting cellular trajectories during differentiation and cell cycle. Additionally, we show how spliced and unspliced integrated features can be used to better classify cells according to sample-associated phenotypes acquired after an experimental perturbation or within a disease state. Lastly, by benchmarking eight data integration methods on the aforementioned prediction tasks, we elucidate method performance specific to gene expression modalities or tasks. While intermediate integration approaches such as SNF, Grassmann joint embedding, integrated diffusion, and PRECISE facilitate increased performance on inferring biological trajectories, simple integration of spliced and unspliced counts through concatenation, sum, or PRECISE achieves increased trajectory inference correlation scores, perturbation classification accuracy, and disease state classification accuracy across most datasets. To this end, integrating multiple gene expression modalities profiled from the same set of cells provides a finer resolution of the transcriptional landscape of development or disease. Thus, an integrated analysis of gene expression modalities may be crucial for the interpretation of dynamic phenotypes.

Several limitations should be considered when integrating gene expression modalities for cellular trajectory inference or disease state classification. In this study, we evaluated methods for constructing integrated graphs or joint embeddings with *a priori* knowledge of ground truth labels. For trajectory inference evaluation, we explored how integrated data influences the change in connectivity or inferred cell state transitions between known cell types identified via FACS. We found that integrated data resulted in increased trajectory inference correlation with respect to a reference trajectory. However, given that the results are sensitive to choice in hyperparameters, it may be challenging to select optimal hyperparameters without *a priori* knowledge of cell types or expected cell type transitions. Here, a range of hyperparameters should be considered when using the intermediate integration methods outlined in this study. Of note, we observed that baseline integration approaches, such as sum and concatenation of spliced and unspliced counts perform consistently well on classifying sample-associated cell phenotypes. This is particularly useful as these approaches are less computationally expensive and do not require hyperparameter tuning. Of note, these baseline methods did not perform well when integrating moments of spliced data with RNA velocity predictions for classification.

Furthermore, the limitations of integration performance are an extension of the modalities used as input. RNA velocity is a noisy extrapolation of gene regulation that can be biased by insufficient sampling of unspliced molecules [57], relies on model assumptions that may be violated [58], and is sensitive to choice in preprocessing tools, such as the quantification of mRNA abundances [59]. Notably, the accuracy of RNA velocity estimation can be improved by incorporating both gene expression and chromatin accessibility data [60]. Moreover, there is currently no consensus on how to appropriately batch effect correct linked gene expression modalities [57]. We chose to jointly correct spliced and unspliced count matrices according to the three metrics and two methods outlined in this study; however, we note that this challenge may bias or limit the interpretation of our results. We anticipate improved performance as bioinformatics tools are developed to better analyze such data. Lastly, although RNA velocity often did not result in an increase in classification accuracy for the datasets selected in this study, this does not preclude it from being informative for the analysis of other datasets. RNA velocity captures gene expression dynamics over the timescale of hours, thus may provide crucial information for longitudinal datasets with finer temporal sampling.

Future work could focus on evaluating temporal gene expression integration for a wider range of tasks, such as unsupervised cell population identification [61], characterizing phenotypic-related cells [40], characterizing differentially abundant cell populations [62, 63], or gene regulatory network inference [64]. This work could also be extended to the analysis of other extrapolated regulatory modalities, including RNA velocity in-situ [14], protein velocity [15], or chromatin velocity [65].

## Methods

### Datasets

We evaluated trajectory inference, experimental perturbation, and disease classification performance on eight datasets spanning various biological contexts. For more details on data preprocessing, see Supplementary Table 1.

#### Hematopoiesis differentiation

FASTQ files consisting of hematopoietic stem and progenitor cells were accessed from Nestorowa et al., [37] with the accession code GSE81682. FACS labels from broad gating were used to annotate six cell populations along three differentiation lineages: long term hematopoietic stem cells (LT-HSC), lymphoid multipotent progenitors (LMPP), multipotent progenitors (MPP), megakaryocyte-erythrocyte progenitors (MEP), common myeloid progenitors (CMP), and granulocyte-monocyte progenitors (GMP) (see Supplementary Table 3). Individual cell FASTQ files were aligned to the mouse reference genome mm10 with the STAR v2.7.7 aligner. A loom file containing spliced and unspliced molecular counts was obtained using Velocyto v0.17.

#### Mouse embryonic cell cycle

A dataset of mouse embryonic stem cells undergoing different stages of the cell cycle was accessed from Buettner et al., [36] with the accession code E-MTAB-2805. FACS cell cycle labels from Hoesct flow sorting were used to annotate cells along three phases: G1, S, and G2/M. Individual cell FASTQ files were aligned to the mouse reference genome mm10 with the STAR v2.7.7 aligner. A loom file containing spliced and unspliced molecular counts was subsequently generated with Velocyto v0.17.

#### LPS stimulation

FASTQ files were accessed from Lane et al., [38] with the accession code GSE94383. Here, a macrophage-like cell line RAW 264.7 was stimulated with lipopolysaccharide (LPS) over 4 time points: 0min unstimulated, 75min-, 150min-, 300min-post LPS stimulation. Files were aligned to the mouse reference genome mm10 with the STAR v2.7.7 aligner. A loom file containing spliced an unspliced molecular counts was generated with Velocyto v0.17. Following preprocessing, batch effect correction was performed on the libraries.

#### INFγ Stimulation

Aligned BAM files of pancreatic islet cell INF*γ* stimulation were accessed from Burkhardt et al., [40] with the accession code GSE161465. This dataset consisted of three donors per stimulation condition (control, INF*γ* stimulated). A loom file containing spliced and unspliced molecular counts was generated for each donor and condition with Velocyto v0.17, then subsequently merged into a single file. Following preprocessing, batch effect correction was performed using the donor labels.

#### AML chemotherapy

To assess disease progression, aligned BAM files of an individual patient with AML undergoing chemotherapy were accessed from Pollyea et al., [5] with the accession code GSE116481. Condition labels consisted of three timepoints: d0 untreated, d2-, d4-post Venotoclax and Azacitidine treatment. A loom file containing spliced and unspliced molecular counts for each timepoint was generated with Velocyto v0.17, then merged into a single file. Following preprocessing, batch effect correction was performed on the condition labels.

#### AML matched diagnosis/relapse

Raw FASTQ files were accessed from Stetson et al., [7] with the accession code GSE126068. In this dataset, PBMCs were collected from 5 patients with AML on the onset of diagnosis and following relapse. FASTQ files were aligned to the human reference genome GRCh38 with the STAR v2.7.7 aligner. A loom file containing spliced and unspliced molecular counts was obtained with Velocyto v0.17. Following preprocessing, batch effect correction was performed using the patient labels.

#### MS case/control

Aligned BAM files were accessed from Schafflick et al., [6] with the accession code GSE138266. Here, two biological samples were collected from each patient (CSF, PBMCs) with a disease status label (control or MS). Loom files containing spliced and unspliced molecular counts for each patient sample were obtained with Velocyto v0.17. Then a merged loom file consisting of control and MS patient cells was generated for each sample independently. Following preprocessing, batch effect correction was performed using the patient labels.

### Preprocessing

#### Quality control, normalization, and highly variable gene selection

All scRNA sequencing datasets were quality control filtered according to read depth and distributions of counts. Following empty droplet and doublet removal, dying cells were removed by ensuring less than 20 percent of total reads were mapped to mitochondrial transcripts. Genes were filtered out if they were expressed in less than five cells or had less than five counts shared between spliced and unspliced matrices. To perform normalization, we estimated size factors for filtered spliced and unspliced count matrices with Scran pooling normalization v1.20.1 [66]. For datasets with an appreciable batch effect, size factors were subsequently scaled according to median normalization of the ratio of average counts between batches with Batchelor v1.8.0; this ensures data is downsampled based upon the batch with the smallest read depth. To restrict the feature space, we selected highly variable genes on log+1 transformed data by estimating a normalized dispersion measure [67] using the highly variable genes function in Scanpy v1.8.1 (flavor = seurat, minimum mean = 0.012, minimum dispersion = 0.25, maximum mean = 5).

#### Batch effect correction

RNA velocity relies on an ordinary differential equation framework to estimate the relationship between two connected modalities, spliced and unspliced mRNA counts [68, 12, 13]. As such, correcting each modality independently may lead to incorrect model fitting and spurious velocity vectors [57]. We evaluated the performance of batch effect correction methods, ComBat [69] and mutual nearest neighbors (MNN) [70] on correcting count data simultaneously. These methods were chosen as they directly correct the original gene expression data. Briefly, we considered two simple approaches for combining the data prior to correction (1) summed spliced and unspliced counts or (2) cell-wise concatenation. To obtain corrected count matrices for summed input data, we followed the batch effect correction approach introduced in in Ref. [71],

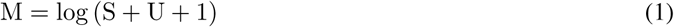

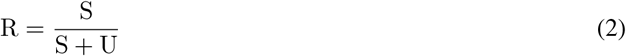

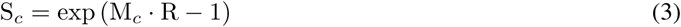

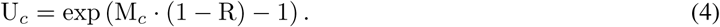

Here, *S* and *U* represent spliced and unspliced count matrices, respectively. Batch effect correction was performed on the summed total expression matrix, *M*, to yield a corrected data matrix M_*c*_. Corrected spliced S_*c*_ and unspliced U_*c*_ counts were then obtained by inverting the log transformation through exponentiation. ComBat was run in python using Scanpy v1.8.1 and MNN was run in R using Batchelor v1.8.0.

#### Batch effect correction evaluation

To evaluate batch effect correction methods on combined spliced and unspliced modalities, we consider three metrics for assessing batch effect removal while preserving both biological variation and the unspliced to spliced relationship.

1. *k-nearest neighbor batch effect correction test (kBET):* The kBET algorithm [72] quantifies batch effects by comparing the batch label composition of local random neighborhoods to the overall global label composition through a *χ*^2^ test. Tests are then averaged to obtain an overall rejection rate. To test for batch effects, we perform kBET using a fixed neighborhood size of *k* = 10 neighbors for each correction approach (uncorrected, MNN sum, MNN concatenation, ComBat sum, ComBat concatenation). kBET scores were computed using kBET v0.99.6.
2. *Local Inverse Simpson’s Index (LISI):* The LISI score [73] measures the degree of batch label mixing by computing the number of cells that can be drawn from a local neighborhood before a batch label is observed twice. Here, local distances are weighted according to a Gaussian kernel and probabilities are determined by the inverse Simpson’s index. LISI returns a diversity score ranging from 1 to the total number of batches. To test for batch label diversity, we compute LISI using a fixed perplexity of 30 for each correction approach (uncorrected, MNN sum, MNN concatenation, ComBat sum, ComBat concatenation). LISI scores were computed using harmonypy.
3. *Pearson correlation of phase space pairwise distances:* The dynamical model of RNA velocity estimates transcriptional dynamics by inferring gene-specific reaction rate and latent parameters through an expectation-maximization framework on the phase space (spliced and unspliced counts) of the data. To quantify how well a batch effect correction approach preserves the unspliced to spliced relationship across all cells, we compared phase space cellular neighborhoods by computing the Pearson correlation of pairwise distances in the phase space for each donor and pairwise distances of the same cells in corrected data. In other words, for each gene we obtain a single correlation score capturing how well cell-cell distances are preserved in the phase space of corrected data with respect to an individual donor/patient. The distribution of gene correlations measure the overall quality of correction for retaining similar cell distributions for RNA velocity fitting and estimation.

To select a batch effect correction approach, we evaluated correction performance on the each biological condition individually. Furthermore, we took the intersection of genes that were highly variable across all profiled samples (e.g. libraries, donors, patients) to ensure that the data being compared were specific to the biological system under study and that donor-specific variation was removed. For each dataset, we selected the batch effect correction approach that had the best performance across all three metrics (see Supplementary Table 1, Supplementary Figure 16). One exception was the AML diagnosis/relapse dataset, which contained too few cells for the analysis. Here, we selected ComBat concatenation, as it was the approach that consistently performed well on all other datasets. Once an approach was selected, we performed joint correction on the original full dataset as outlined previously (See Preprocessing).

#### RNA velocity estimation

To estimate RNA velocity, we used the dynamical model implementation in Scvelo v0.2.3. More specifically, first order moments of spliced and unspliced counts were computed based on a *k*-nearest neighbor graph of cells (*k* = 10), constructed by calculating pairwise Euclidean distances between cells based on their first 50 principal components (PCs). The full dynamical model was then solved for all genes to obtain a high dimensional velocity vector for every cell. Given that populations of cells may have different mRNA splicing and degradation kinetics, we performed a likelihood ratio test for differential kinetics on the clusters identified from Leiden community detection (resolution parameter of 1.0) [74]. Clusters of cells that exhibited different kinetic regimes were fit independently and velocity vectors were corrected.

#### Sketching

To evaluate integration performance on the large-scale case/control datasets, we first performed subsampling with geometric sketching. Geometric sketching [75] is an algorithm that aims to select a representative subset of cells that preserves the overall transcriptional heterogeneity of the full dataset. By approximating the underlying geometry of the data through a plaid covering of equal volume hypercubes, geometric sketching is able to evenly select cells such that rare cell types are sufficiently sampled. We implemented geometric sketching to select a representative subset of cells from both Multiple Sclerosis case/control datasets. Sketches were constructed from the transcriptional landscape of the mature gene expression data (spliced or moments of spliced), with sketch sizes of approximately twenty percent of the total data. Sketch indices were then used to subsample all modalities prior to integration and disease state classification evaluation.

### Integration methods

#### Problem Formulation

Let 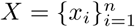 denote a single-cell dataset consisting of one gene expression modality, where *x*_*i*_ ∈ ℝ^*d*^ represents a vector of *d* genes measured in cell *i*. Given a collection of *m* gene expression modalities 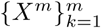 sampled from *N* individuals, where for sample *i* there is an associated label *y*_*i*_, our goal is to identify a biologically meaningful consensus representation, 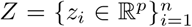 where *p* represents shared latent features such that *p* ≤ *d*. In this case, we wish to use this consensus representation to build a predictive model to infer biological trajectories or to predict the patient-specific or treatment-induced phenotypic label for sample *i, y*_*i*_. In this section, we describe the methods selected for integrating two groups of gene expression modalities, either moments of spliced counts with RNA velocity data or normalized and log transformed spliced and unspliced count matrices. For more details on implementation and hyperparameter tuning, see Supplementary Table 2.

##### Unintegrated

To evaluate a baseline approach representing unintegrated data, we constructed a *k*-nearest neighbor graph (*k* = 10) from the top 50 principal components, generated from the normalized and log transformed spliced counts. This is akin to what is traditionally used for downstream single-cell analysis, as outlined by current best practices [48].

##### Concatenation

Gene expression data matrices were horizontally concatenated to obtain a merged data matrix with dimensions *n* × 2*d*. Principal Component Analysis (PCA) was performed on the concatenated matrix and a *k*-nearest neighbors graph (*k* = 10) of cells was ultimately constructed based on the top 50 principal components.

##### Sum

Gene expression data matrices were summed to obtain a merged data matrix with dimensions *n* × *d*. PCA was performed on the summed matrix and a *k*-nearest neighbor graph (*k* = 10) was constructed from the top 50 principal components.

##### CellRank

CellRank [17] computes a joint transition probability matrix through a weighted sum of expression and velocity transition probability matrices as,

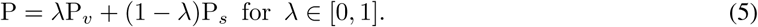

Here, P_*v*_ represents the velocity transition matrix, P_*s*_ represents the expression similarity transition matrix, and *λ* is the weight parameter. We used CellRank v1.1.0 and performed hyperparameter tuning by varying the weight parameter *λ*, the measure of velocity similarity (correlation, dot product, or cosine), and the model that determines if velocity uncertainty is propagated in the transition matrix computation (monte-carlo, dynamical). Given that this approach relies on RNA velocity directionality, integration was only performed using moments of spliced and RNA velocity data.

##### PRECISE

PRECISE [47] was adapted to integrate temporal gene expression modalities. PRECISE first computes principal components for each modality individually, then geometrically aligns components to extract common principal vectors that represent similar weighted combinations of genes. From here, a consensus feature representation is computed by optimizing the match between interpolated sets of features (e.g. expression and velocity). For this analysis, we obtained a lower dimensional latent space by projecting expression data onto (1) the principal vectors (denoted as PRECISE) or (2) the consensus features (denoted as PRECISE consensus). From this shared embedding space, we constructed a *k*-nearest neighbor graph (*k* = 10). For both approaches, we performed hyperparameter tuning by varying the number of included principal vectors. Given that the principal vectors are rank ordered according to modality similarity, selection is analogous to filtering the data based on shared or unshared information. PRECISE v1.2 was used and modified to include dissimilar components.

##### Similarity Network Fusion

Similarity Network Fusion (SNF) [25] constructs a joint graph of cells according to gene expression data modalities using a two step process. First a cell affinity graph 𝒢^*m*^ = (𝒱^*m*^, *ε*^*m*^) is computed for each modality, where 𝒱^*m*^ represents cells and edges, *ε*^*m*^, are weighted according to modality-specific similarity using a heat kernel as follows. Here, we compute 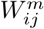, which gives the specific edge-weight between cells *i* and *j* in modality *m* as,

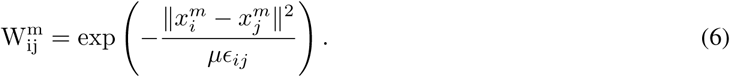

Specifically, *W*^*m*^ is a *n* × *n* similarity matrix for modality *m, μ* is a scaling hyperparameter, and *ϵ*_*ij*_ is a bandwidth parameter that takes into account local neighborhood sizes. Next, the two individual modality networks are integrated through nonlinear diffusion iterations between each modality to obtain a fused network. Importantly, the network fusion step ensures that the merged graph representation retains edge similarities that are strongly supported by an individual modality in addition to similarities shared across modalities. To compare results to the intermediate embedding integration methods, we modified SNF by constructing a shared embedding from the fused network through eigendecomposition of the unnormalized graph Laplacian L_*u*_. Note that L_*u*_ is computed as,

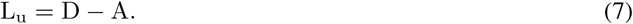

Here, *D* is a diagonal degree matrix with *i*-th diagonal element, *d*_*i*_ = ∑_*j*_ *A*_*ij*_ and *A* is the symmetric merged SNF affinity adjacency matrix. Given that eigenvectors of the Laplacian represent frequency harmonics, we selected the eigenvectors corresponding to the *K* smallest eigenvalues to low pass filter high frequency noise [76]. We then constructed a *k*-nearest neighbor graph (*k* = 10) for evaluation. We performed hyperparameter tuning by varying the number of nearest neighbors, the bandwidth scaling parameter *μ*, and the number of eigenvectors for the merged graph embedding. SNF was implemented using the snfpy v0.2.2 package in python.

##### Grassmann Joint Embedding

The Grassmann joint embedding approach introduced in Ref. [26] was adapted to construct a shared representative subspace of temporal gene expression information. Similar to SNF, the Grassmann embedding approach begins by constructing affinity matrices to encode similarities between cells *i* and *j* in each modality using a heat kernel as,

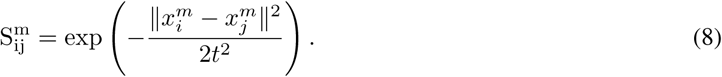

Here, *S*^*m*^ is a *n* × *n* between-cell similarity matrix for modality *m* and *t* is the kernel bandwidth parameter. To prioritize local similarities, the *k*-nearest neighbors according to the similarity matrix *S*^*m*^ are identified and the similarity matrix is further redefined as,

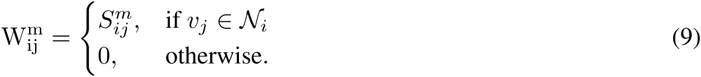

Here, cell *v*_*i*_ and *v*_*j*_ are connected with an edge with edge weight *S*_*ij*_ if the cell is within the set of *v*_*i*_’s neighbors, 𝒩_*i*_. Next, low-dimensional subspaces are computed through eigendecomposition of the normalized graph Laplacian of each data type. The normalized graph Laplacian 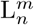, is formally defined as,

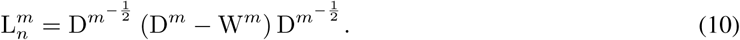

Here, *m* indexes the data modality and D^*m*^ represents a diagonal degree matrix, such that the *i*-th diagonal element, 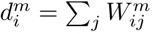. Furthermore, A^*m*^ is the symmetric Grassmann affinity matrix of modality *m*. A shared representative subspace from [26] is then ultimately computed as,

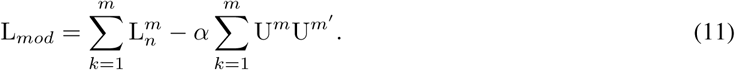

Here, U^*m*^ represents an individual subspace representation and *α* controls the trade-off between preserving modality-specific structural similarities (in the first term) and minimizing the distance between each subspace representation (in the second term). Lastly, an eigendecomposition of the Laplacian of the joint graph L_*mod*_ was computed to extract the *K* eigenvectors corresponding to the first *K* eigenvalues to represent the merged embedding space. For evaluation, we constructed a *k*-nearest neighbor graph (*k* = 10) from this shared space. Hyperparameter tuning was performed by varying the number of nearest neighbors and kernel bandwidth parameter *t* in the affinity graph construction, as well as *α*, and the number of eigenvectors to include for the merged graph embedding.

##### Integrated Diffusion

Integrated diffusion [24] combines data modalities by computing a joint data diffusion operator. First, individual modalities are locally denoised by performing a truncated singular value decomposition (SVD) on local neighborhoods determined through spectral clustering. Next a symmetric diffusion operator is constructed for each denoised modality, and spectral entropy is used to determine the number of diffusion time steps to take. By taking the reduced ratio of information, the joint diffusion operator P_*j*_ is computed as,

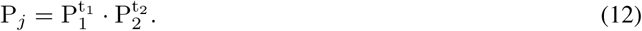

Here, P_1_ and P_2_ represent individual modality diffusion operators (e.g. expression and velocity) and t_1_ and t_2_ represent the reduced ratio of diffusion time steps, respectively. By powering transition probability matrices independently, this captures both modality-specific information, while allowing the random walk to jump between data types for merging. Lastly, the joint diffusion operator is powered using the same spectral entropy measure. It is important to note that choice of *t* is crucial, as it can either effectively denoise data or remove important variation and lead to oversmoothing. We eigendecomposed the diffused joint operator and selected the eigenvectors corresponding to the *K* largest eigenvalues to obtain a merged lower dimensional representation. We then constructed a *k*-nearest neighbor graph (*k* = 10). Hyperparameter tuning was performed by varying the number of clusters for local denoising, the number of nearest neighbors in affinity graph construction, and the number of included eigenvectors.

### Evaluation

#### Trajectory inference

To quantify how well incorporation of unspliced counts or RNA velocity recapitulates the underlying biological trajectory, we compared predicted trajectories to a ground truth reference using the metrics implemented in the R suite Dynverse [51]. Reference trajectories were curated from the literature [37, 36, 54], with cell groups, connections, and root cluster provided by the authors of the original study. We note that cell population annotations were externally determined through cell surface protein measurements and not from unsupervised clustering on the expression data.

To obtain predicted trajectories from integrated data, we performed trajectory inference using Partition-based Graph Abstraction [52] followed by diffusion pseudotime [53], as this approach was shown to outperform other methods for inference of complex differentiation topologies [51]. Predicted trajectories consisted of two main attributes: (1) a trajectory network, where nodes represent FACS cell groups and edges connect populations based on PAGA inferred connectivity and (2) a list of cellular percentages representing a cell’s relative position between groups. Here, cellular percentages were determined from diffusion pseudotime using 20 diffusion map components generated from the underlying integrated or unintegrated *k*-nearest neighbor graph. For each integration approach, we computed predicted trajectories for ten random root cells selected from the annotated root cluster.

To evaluate a method’s performance on inferring developmental gene expression dynamics from integrated or unintegrated data, we compared reference and predicted trajectories using two metrics previously described in Ref. [51]: cell distance correlation and feature importance score correlation.

1. *Cell distance correlation* C_*corr*_: Geodesic distances represent the shortest path distance between two cells on a nearest neighbor graph of the data [77]. To estimate a measure of the correlation of between-cell distances between reference and predicted trajectories, geodesic distances were computed between cells on a trajectory graph. The cell distance correlation is defined as the Spearman rank correlation between the geodesic cell distances of both trajectories.
2. *Feature importance score correlation* F_*corr*_: To assess whether the same temporally expressed genes were found in the predicted trajectory as in the reference, a random forest regression framework was used to predict the expression values of each gene based on geodesic distances of each cell to each cell state cluster. The feature importance score correlation is defined as the Pearson correlation between the reference and predicted scores.

To obtain an overall trajectory inference correlation score reflective of high cell and feature similarity, we compute the harmonic mean of both correlation metrics as,

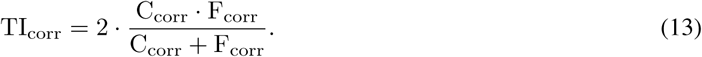

### Classification

#### Label Propagation

To quantitatively compare integration methods on disease state prediction, we aimed to implement an approach that would use the underlying integrated or unintegrated graph structure. Label propagation [56] is a semi-supervised learning algorithm that uses iterative diffusion processes to predict the labels of unlabeled nodes. The output of this algorithm is a probability distribution of labels for every cell. We implemented label propagation to predict stimulation condition or disease status labels as follows.

Let 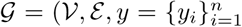 denote a graph of *n* cells comprising the nodes (𝒱) generated from an integration approach and the set *ε* edges encoding between-cell similarities. Similarly, a particular *y*_*i*_ gives a phenotypic label for cell *i* (e.g. patient disease status). Let *y′* = (*y*_*l*_, *y*_*u*_) denote a vector consisting of a training subset of cells that are labeled 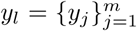 where *y*_*j*_ ∈ *y* and *m* < *n*, and a test subset of cells that are unlabeled, *y*_*u*_ = {0}^*n*−*m*^. Given 𝒢 and *y′*, our goal is to assign a label to the unlabeled cells and the corresponding entries of *y′s*. To do so, we perform the following approach.

1. Stratified random sampling is used to assign cells to a training or test set; this ensures that the original ratio of class labels (e.g. AML diagnosis or relapse) remains the same as in the full dataset.
2. Initialize algorithm on the training set to predict the labels of the masked test set. Each node has a label 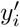, and edge weight *w*_*ij*_ representing the strength of similarity between nodes *i* and *j*. Here, larger weights indicate a higher probability of cell *i* propagating its label 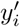 to cell *j*.
3. Labels are iteratively updated through diffusion, where D is a diagonal degree matrix with *i*’th diagonal element *d*_*i*_ = ∑_*j*_ *W*_*ij*_ as,

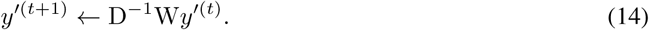
4. Row normalize labels *y′* to maintain a probability distribution.
5. Training labels are clamped after each iteration as,

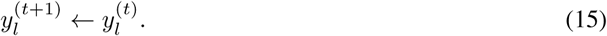
6. Iterations are repeated until convergence, with a threshold *δ* = 0.001, such that,

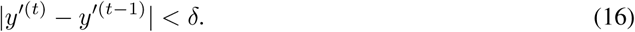
7. Class labels are assigned to every node by taking the label with the maximum probability.

We repeated this procedure for ten random training initializations to obtain a set of predicted labels for each integration approach.

#### Support Vector Machine (SVM)

The support vector machine (SVM) [78] is a supervised learning algorithm that constructs hyperplanes in the high dimensional data to separate classes. We implemented SVM as a secondary classification approach for predicting perturbation response or disease status labels from the individual or joint embedding space (e.g. PCA, diffusion embedding). Specifically, nested 10-fold cross validation was performed using stratified random sampling to assign cells to either a training or test set. SVM hyperparameters were tuned over a grid search within each fold prior to training the model and labels were subsequently predicted from the test data.

#### Metrics

To quantify stimulation condition and disease status classification performance, we compared predicted labels to ground truth annotations using three metrics: F1 score, balanced accuracy (acc_b_), and area under the receiver operator curve (AUC). The F1 score measures the harmonic mean of precision and recall as,

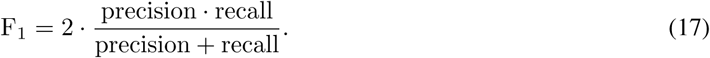

Balanced accuracy represents the average of sensitivity (true positive rate) and specificity (true negative rate). When predicting more than two labels (e.g. disease progression), we computed the mean sensitivity for all classes.

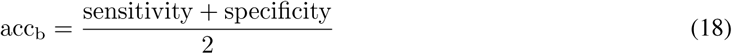

Lastly, area under the receiver operator curve was computed using the soft probability assignments. For the multi-class case, each class label was compared to the remaining in an all vs. rest approach, then averaged. All of these metrics return a value between 0 and 1, where 1 indicates predicted labels were in perfect accordance to the ground truth annotations.

### Aggregate scores

To rank methods for each prediction task, we compute aggregate scores by taking the mean of scaled method scores across datasets. More specifically, we first define an overall method score per dataset as the median of each metric. Method scores were subsequently min-max scaled to ensure datasets were equally weighted prior to computing the average.

## Data and code availability

The raw publicly available single-cell RNA sequencing datasets downloaded and used in this study are available in the Gene Expression Omnibus repository, under the accession codes GSE81682 for hematopoiesis differentiation [37], GSE94383 for LPS stimulation [38], GSE161465 for INF*γ* stimulation [40], GSE11648 for AML chemotherapy [5], GSE1260681 for AML diagnosis/relapse [7], and GSE138266 for MS case/control PBMC and CSF datasets [6] and in the European Nucleotide Archive repository, under accession numbers E-MTAB-2805 for mouse embryonic cell cycle [36] datasets, respectively. Loom files and preprocessed data are available in the Zenodo repository https://doi.org/10.5281/zenodo.6110279. All functions for preprocessing, integration, and evaluation are available at www.github.com/jranek/EVI.

## Authors’ contributions

JSR, NS, JEP conceptualized and designed the study. JSR performed data preprocessing, benchmarking, evaluation, and analysis. JSR wrote the manuscript with input from all authors. All authors read and approved of the final manuscript.

## Acknowledgements

The authors would like to thank Logan Whitehouse and Tarek Zikry for their thoughtful discussions related to this work.

## Supplementary Tables

**Supplementary Table 1:** Datasets and preprocessing overview

| ID | Description | Metadata | Task | Platform | Organism | Reference | Batch | Batch correction approach | Normalization |
| --- | --- | --- | --- | --- | --- | --- | --- | --- | --- |
| Nestorowa | hematopoiesis differentiation | FACS | TI | Smart-seq2 | Mouse | mm10 | NA | NA | Scran |
| Buettner | mouse embryonic cell cycle | FACS | TI | Smarter C1 | Mouse | mm10 | NA | NA | Scran |
| Lane | LPS stimulation | condition | classification | Smart-seq2 | Mouse | mm10 | library | ComBat concatenation | Scran with batch |
| Pollyea | AML chemotherapy | condition | classification | 10X Genomics | Human | GRCh38 | condition | ComBat concatenation | Scran with batch |
| Burkhardt | IFN-γ stimulation | condition | classification | 10X Genomics | Human | GRCh38 | patient | MNN concatenation | Scran with batch |
| Stetson | AML diagnosis/relapse | disease status | classification | Smart-seq2 | Human | GRCh38 | patient | ComBat concatenation | Scran with batch |
| Schafflick | MS case/control | disease status | classification | 10X Genomics | Human | GRCh38 | patient | ComBat concatenation | Scran with batch |

**Supplementary Table 2:**
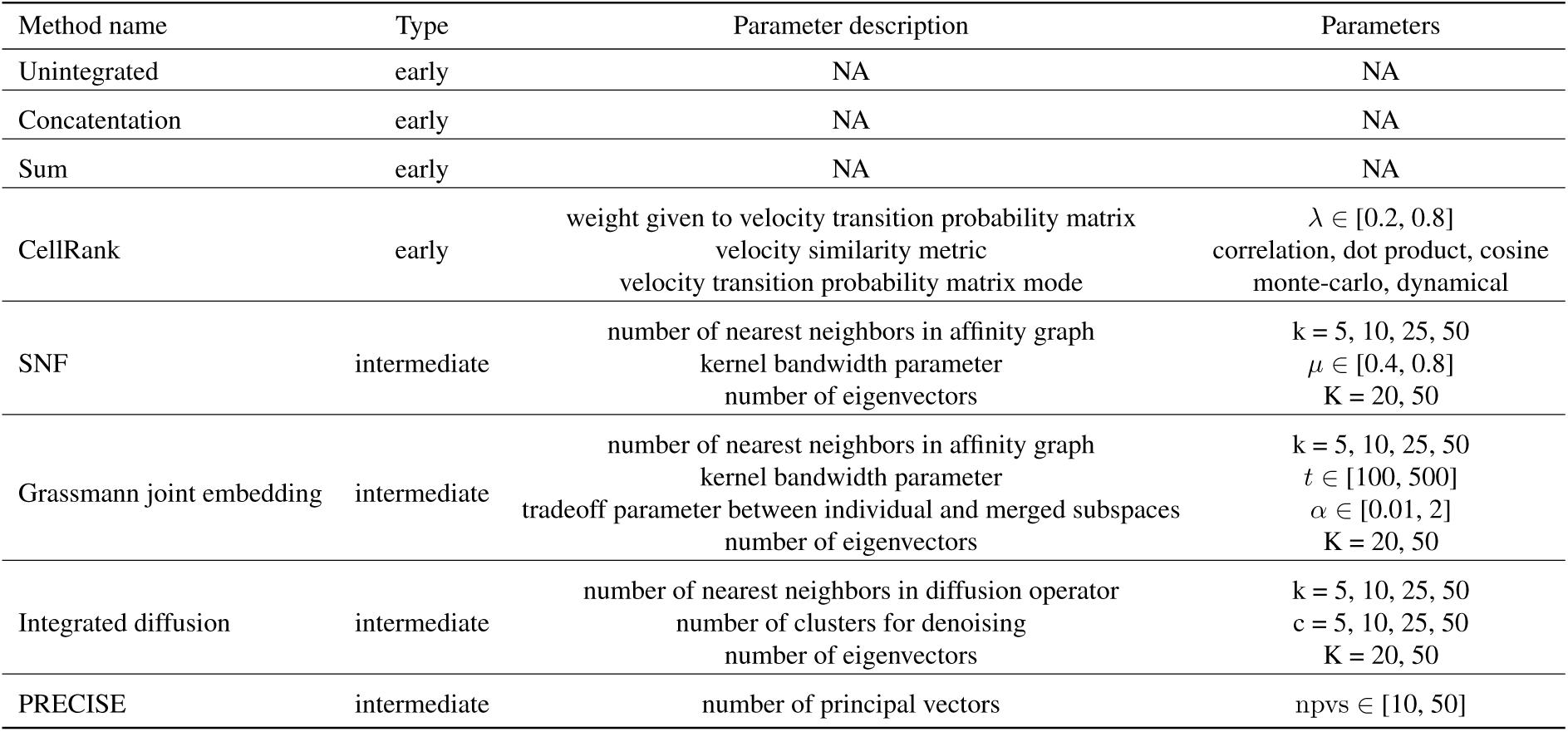
Overview of optimized parameters

**Supplementary Table 3:** Surface markers for hematopoietic stem and progenitor cells in Nestorowa et al.

| name | ID | markers |
| --- | --- | --- |
| long-term hematopoietic stem cells | LT-HSC | $\text{Lin}^- \text{c-Kit}^+ \text{Sca1}^+ \text{Flk2}^- \text{CD34}^-$ |
| lymphoid multipotent progenitors | LMPP | $\text{Lin}^- \text{c-Kit}^+ \text{Sca1}^+ \text{Flk2}^+ \text{CD34}^+$ |
| multipotent progenitors | MPP | $\text{Lin}^- \text{c-Kit}^+ \text{Sca1}^+ \text{Flk2}^- \text{CD34}^+$ |
| megakaryocyte-erythrocyte progenitors | MEP | $\text{Lin}^- \text{c-Kit}^+ \text{Sca1}^- \text{CD16/32}^- \text{CD34}^-$ |
| common myeloid progenitors | CMP | $\text{Lin}^- \text{c-Kit}^+ \text{Sca1}^- \text{CD16/32}^- \text{CD34}^+$ |
| granulocyte-monocyte progenitors | GMP | $\text{Lin}^- \text{c-Kit}^+ \text{Sca1}^- \text{CD16/32}^+ \text{CD34}^+$ |

## Supplementary Figures

**Supplementary Figure 1:**
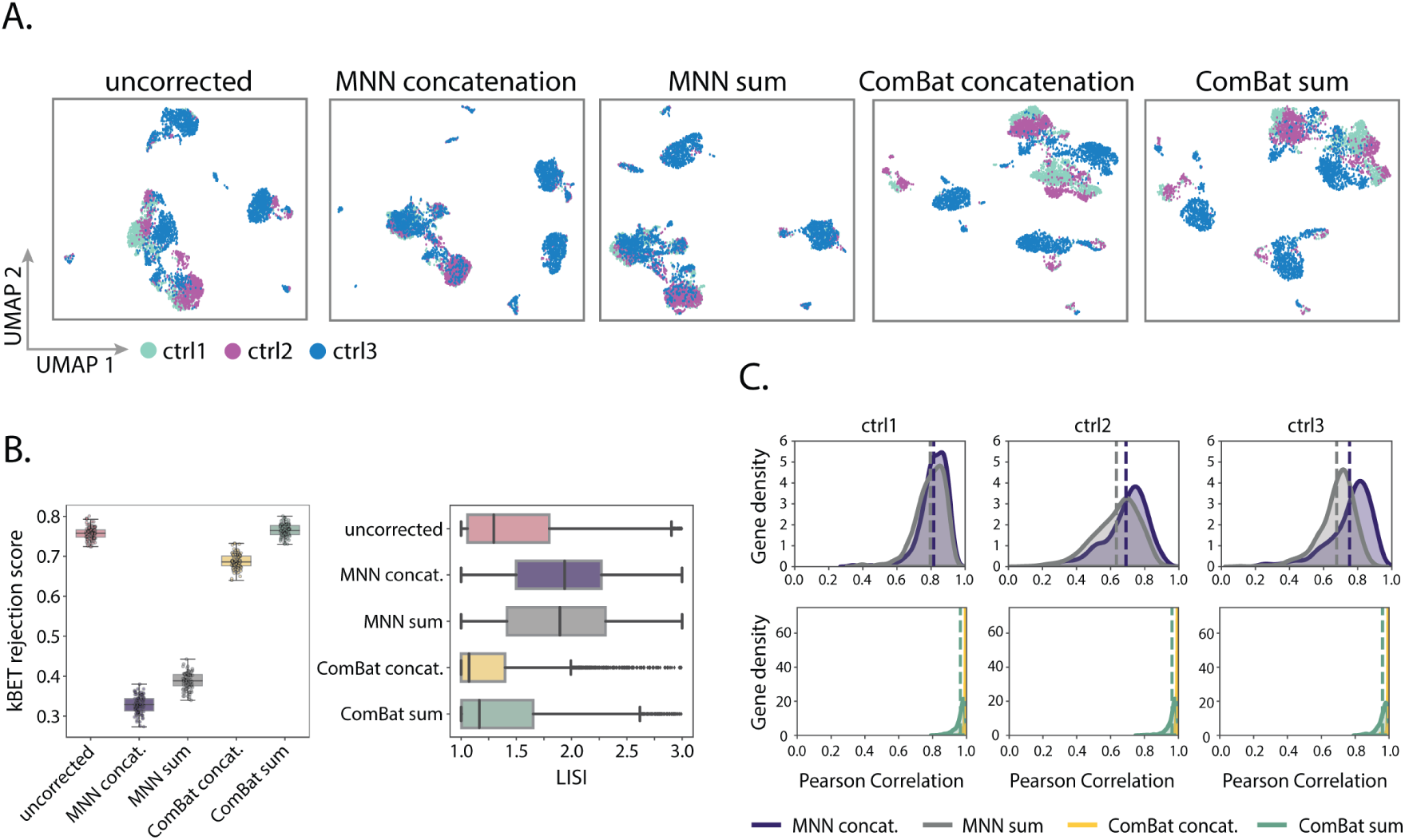
Evaluating batch effect correction for control pancreatic islet cells in INF*γ* stimulation dataset. (A) UMAP visualization of control pancreatic islet cells across batch correction strategies. Spliced and unspliced modalities were combined via concatenation or sum prior to correction with mutual nearest neighbors (MNN) or ComBat. Method performance was measured by batch label mixing metrics kBET and LISI (B), as well as the preservation of the relationship between spliced and unspliced counts (C). Distributions represent the per gene Pearson correlation between cell-cell distances in the phase space (unspliced, spliced) of corrected data and the cell-cell distances in the phase space of each individual donor. Top panel: Pearson correlation of MNN concatenation or MNN sum to control donors. Bottom panel: Pearson correlation of ComBat concatenation or ComBat sum to control donors. Dashed line represents the mean correlation.

**Supplementary Figure 2:**
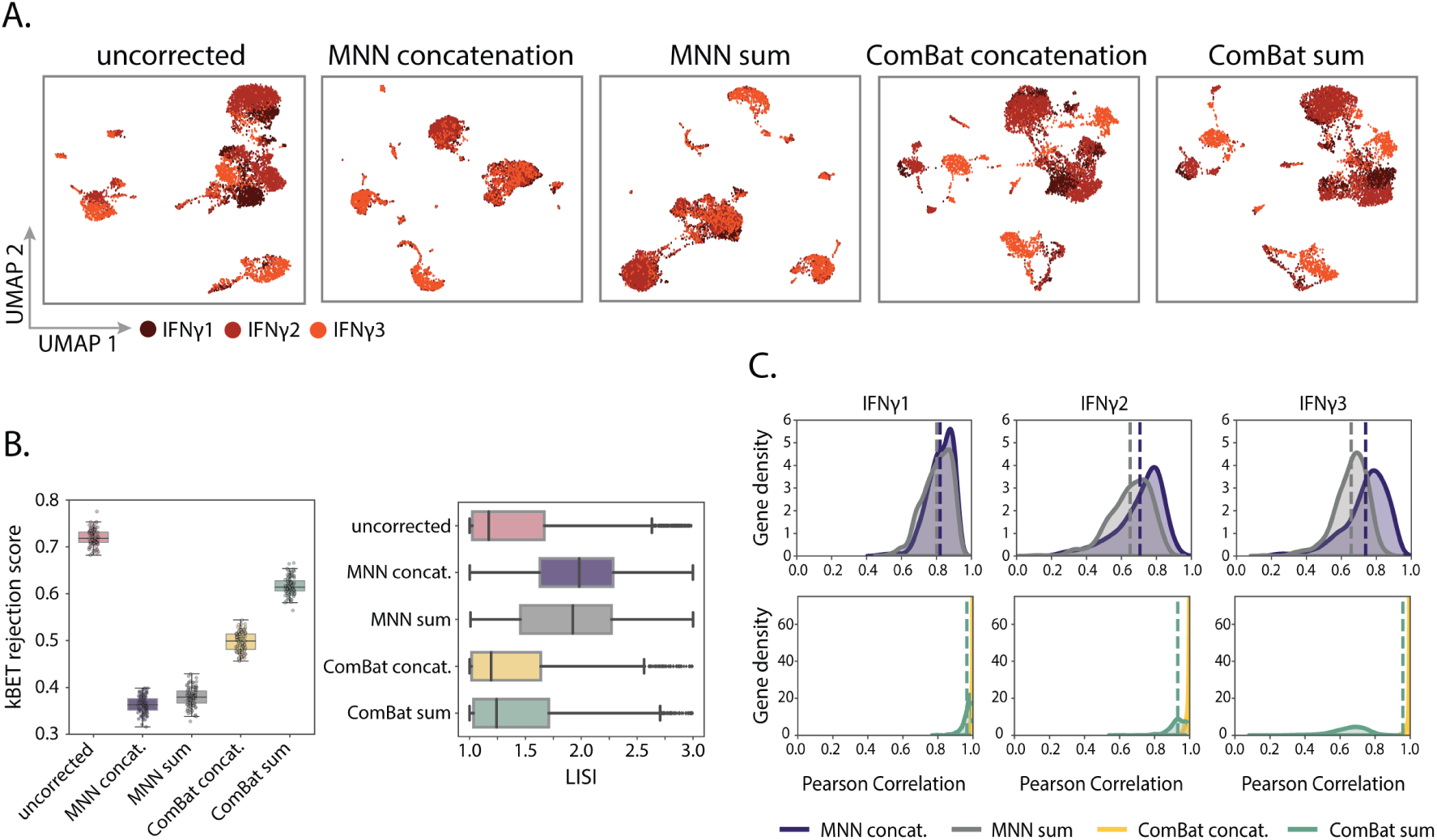
Evaluating batch effect correction for INF*γ* stimulated pancreatic islet cells in INF*γ* stimulation dataset. (A) UMAP visualization of INF*γ* stimulated pancreatic islet cells across batch correction strategies. Spliced and unspliced modalities were combined via concatenation or sum prior to correction with mutual nearest neighbors (MNN) or ComBat. Method performance was measured by batch label mixing metrics kBET and LISI (B), as well as the preservation of the relationship between spliced and unspliced counts (C). Distributions represent the per gene Pearson correlation between cell-cell distances in the phase space (unspliced, spliced) of corrected data and the cell-cell distances in the phase space of each individual donor. Top panel: Pearson correlation of MNN concatenation or MNN sum to INF*γ* stimulated donors. Bottom panel: Pearson correlation of ComBat concatenation or ComBat sum to INF*γ* stimulated donors. Dashed line represents the mean correlation.

**Supplementary Figure 3:**
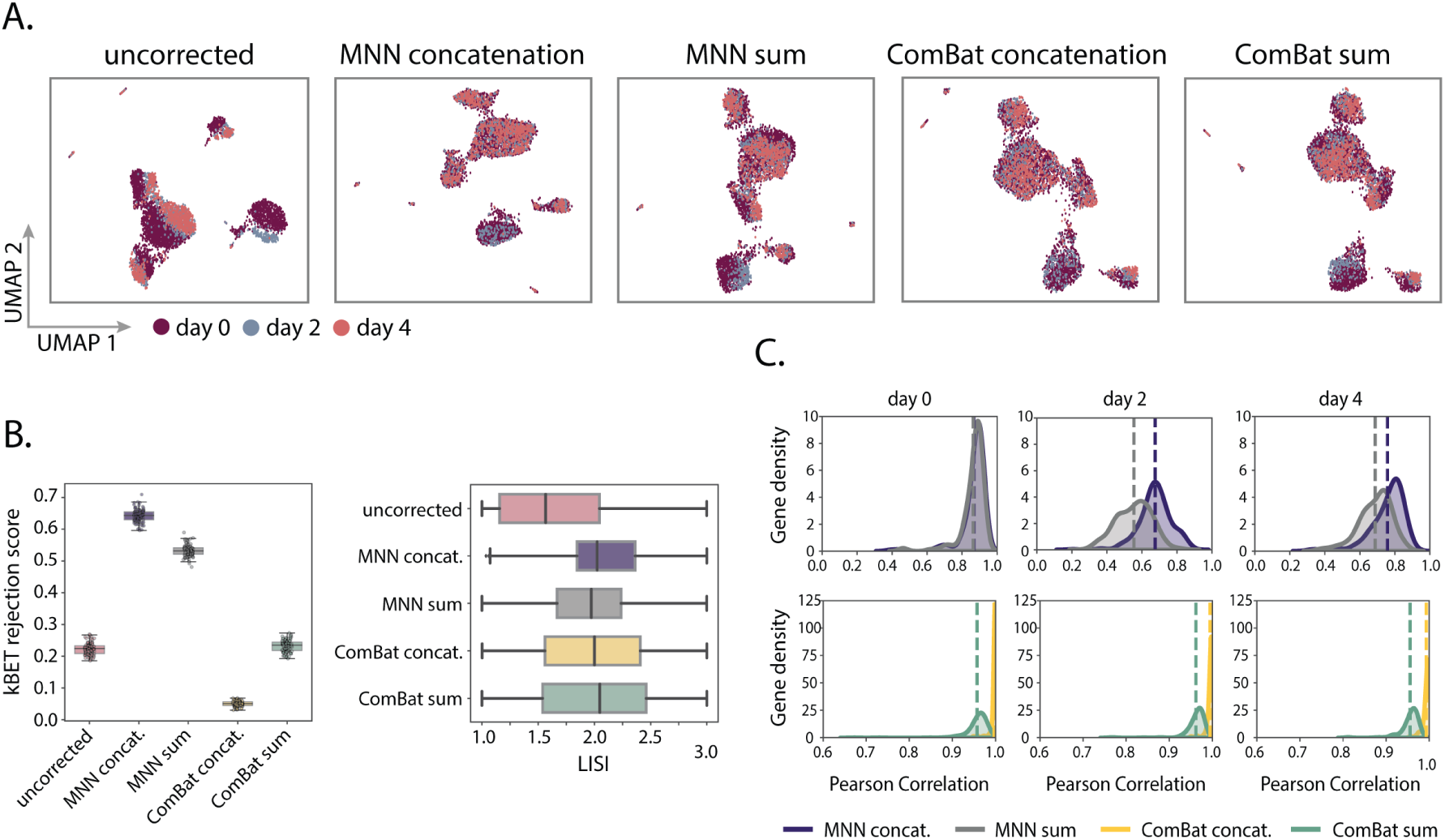
Evaluating batch effect correction for AML chemotherapy treated cells. (A) UMAP visualization of AML chemotherapy treated cells across batch correction strategies. Spliced and unspliced modalities were combined via concatenation or sum prior to correction with mutual nearest neighbors (MNN) or ComBat. Method performance was measured by batch label mixing metrics kBET and LISI (B), as well as the preservation of the relationship between spliced and unspliced counts (C). Distributions represent the per gene Pearson correlation between cell-cell distances in the phase space (unspliced, spliced) of corrected data and the cell-cell distances in the phase space of each time point (d0, d2, d4). Top panel: Pearson correlation of MNN concatenation or MNN sum to individual timepoint. Bottom panel: Pearson correlation of ComBat concatenation or ComBat sum to individual timepoint. Dashed line represents the mean correlation.

**Supplementary Figure 4:**
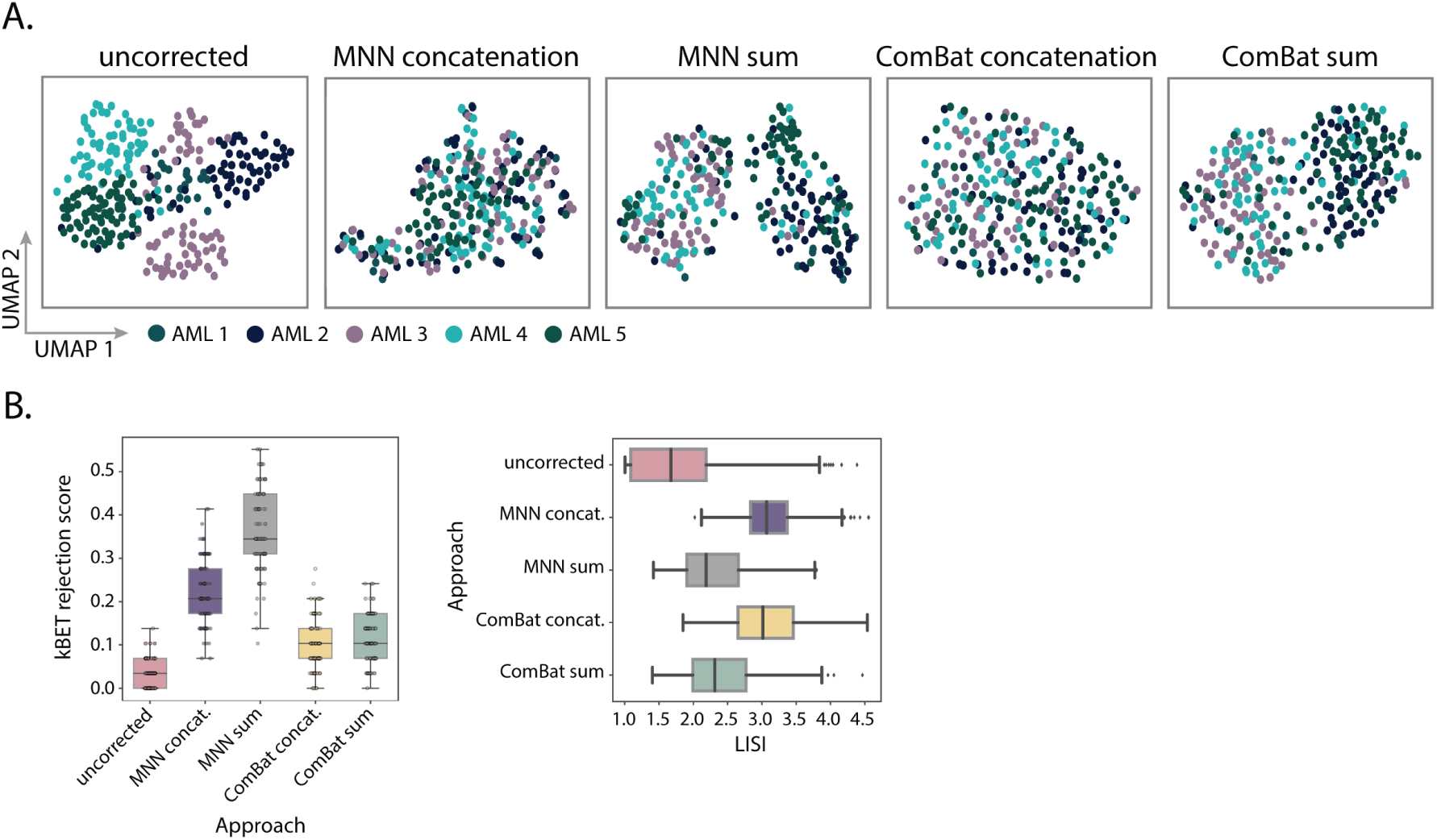
Evaluating batch effect correction for AML diagnosis patient cells in AML diagnosis/relapse dataset. (A) UMAP visualization of AML diagnosis patient cells across batch correction strategies. Spliced and unspliced modalities were combined via concatenation or sum prior to correction with mutual nearest neighbors (MNN) or ComBat. (B) Method performance was measured by batch label mixing metrics kBET and LISI across patients. Dashed line represents the mean correlation.

**Supplementary Figure 5:**
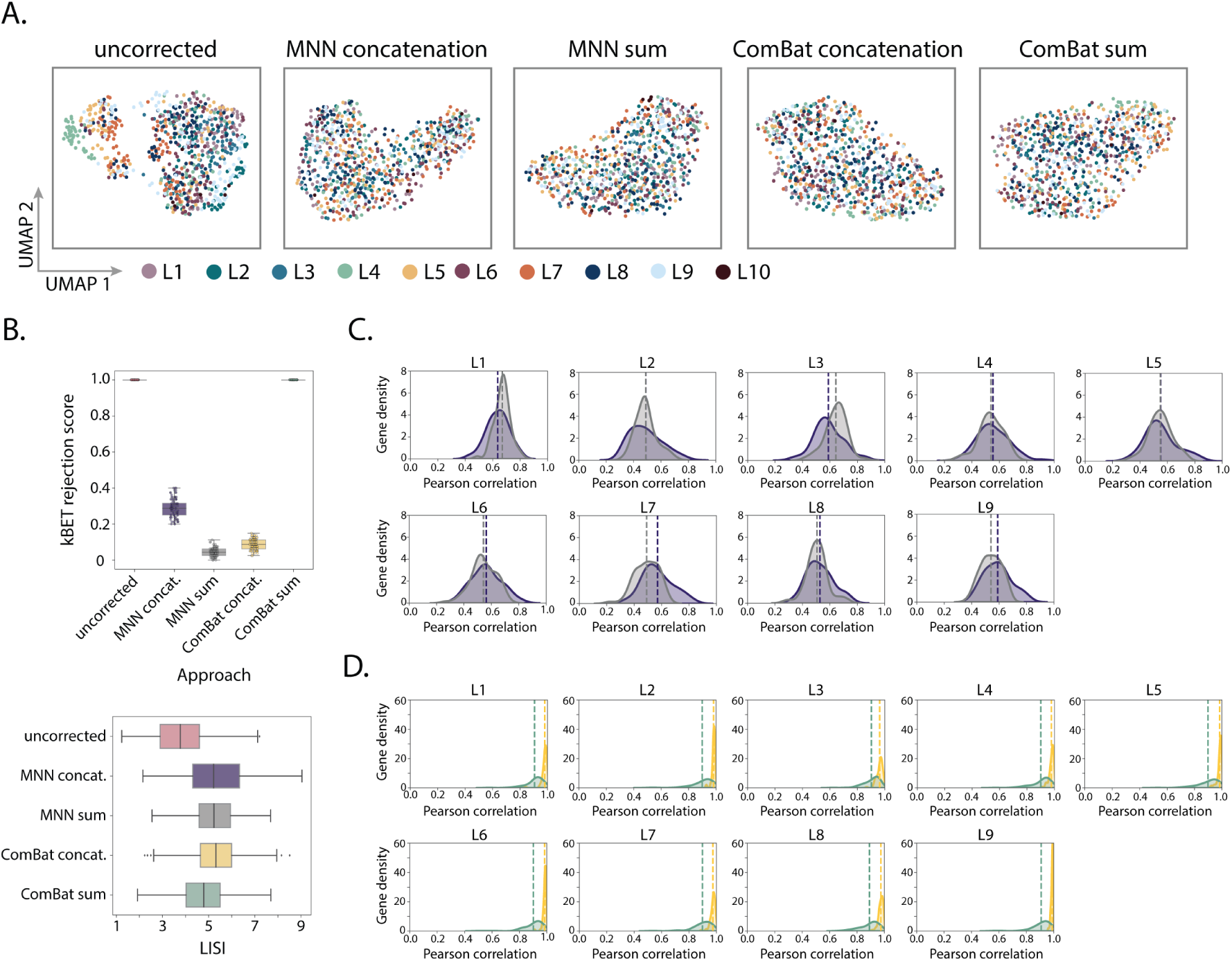
Evaluating batch effect correction for LPS stimulated cells. (A) UMAP visualization of LPS stimulated cells across batch correction strategies. Spliced and unspliced modalities were combined via concatenation or sum prior to correction with mutual nearest neighbors (MNN) or ComBat. Method performance was measured by batch label mixing metrics kBET and LISI (B), as well as the preservation of the relationship between spliced and unspliced counts (C, D). Distributions represent the per gene Pearson correlation between cell-cell distances in the phase space (unspliced, spliced) of corrected data and the cell-cell distances in the phase space of each individual library. Panel C: Pearson correlation of MNN concatenation or MNN sum to individual library. Panel D: Pearson correlation of ComBat concatenation or ComBat sum to individual library. Dashed line represents the mean correlation. Library 10 was excluded from correlation analysis as it contained too few cells.

**Supplementary Figure 6:**
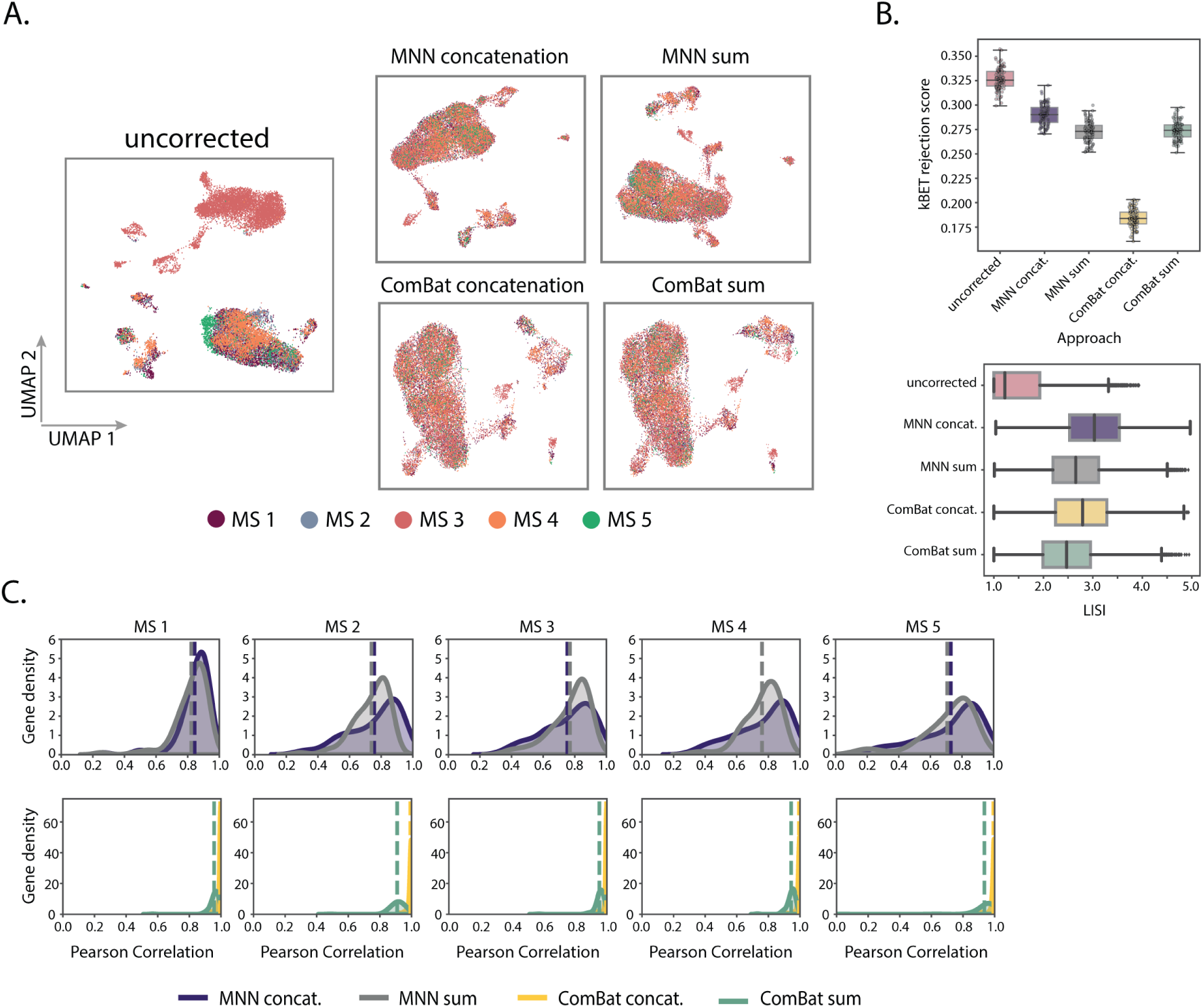
Evaluating batch effect correction for MS patient CSF cells in MS case/control dataset. (A) UMAP visualization of MS patient CSF cells across batch correction strategies. Spliced and unspliced modalities were combined via concatenation or sum prior to correction with mutual nearest neighbors (MNN) or ComBat. Method performance was measured by batch label mixing metrics kBET and LISI (B), as well as the preservation of the relationship between spliced and unspliced counts (C). Distributions represent the per gene Pearson correlation between cell-cell distances in the phase space (unspliced, spliced) of corrected data and the cell-cell distances in the phase space of each individual MS patient. Top panel: Pearson correlation of MNN concatenation or MNN sum to MS patients. Bottom panel: Pearson correlation of ComBat concatenation or ComBat sum to MS patients. Dashed line represents the mean correlation.

**Supplementary Figure 7:**
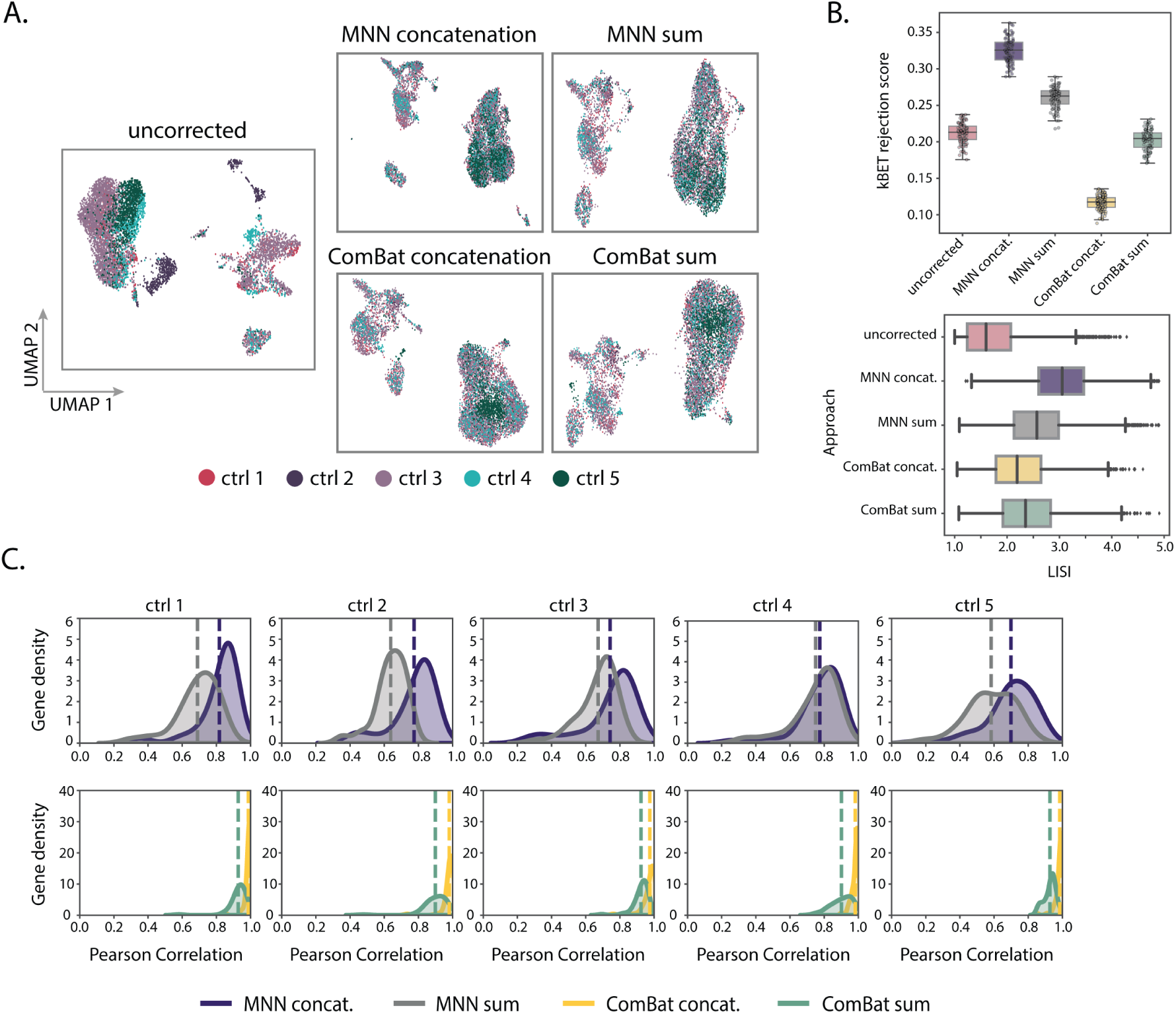
Evaluating batch effect correction for control patient CSF cells in MS case/control dataset. (A) UMAP visualization of control patient CSF cells across batch correction strategies. Spliced and unspliced modalities were combined via concatenation or sum prior to correction with mutual nearest neighbors (MNN) or ComBat. Method performance was measured by batch label mixing metrics kBET and LISI (B), as well as the preservation of the relationship between spliced and unspliced counts (C). Distributions represent the per gene Pearson correlation between cell-cell distances in the phase space (unspliced, spliced) of corrected data and the cell-cell distances in the phase space of each individual control patient. Top panel: Pearson correlation of MNN concatenation or MNN sum to control patients. Bottom panel: Pearson correlation of ComBat concatenation or ComBat sum to control patients. Dashed line represents the mean correlation.

**Supplementary Figure 8:**
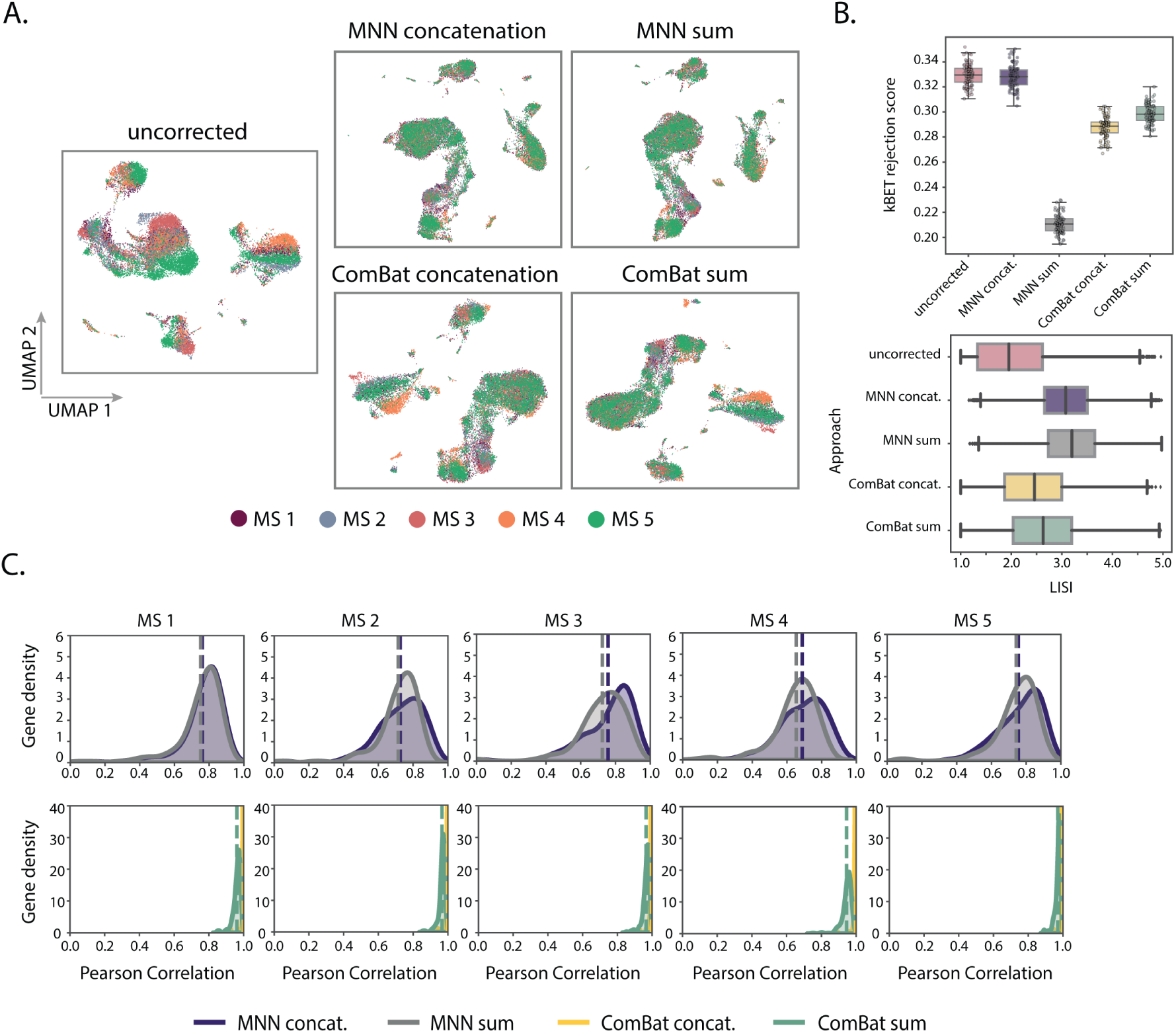
Evaluating batch effect correction for MS patient PBMCs in MS case/control dataset. (A) UMAP visualization of MS patient PBMCs across batch correction strategies. Spliced and unspliced modalities were combined via concatenation or sum prior to correction with mutual nearest neighbors (MNN) or ComBat. Method performance was measured by batch label mixing metrics kBET and LISI (B), as well as the preservation of the relationship between spliced and unspliced counts (C). Distributions represent the per gene Pearson correlation between cell-cell distances in the phase space (unspliced, spliced) of corrected data and the cell-cell distances in the phase space of each individual MS patient. Top panel: Pearson correlation of MNN concatenation or MNN sum to MS patients. Bottom panel: Pearson correlation of ComBat concatenation or ComBat sum to MS patients. Dashed line represents the mean correlation.

**Supplementary Figure 9:**
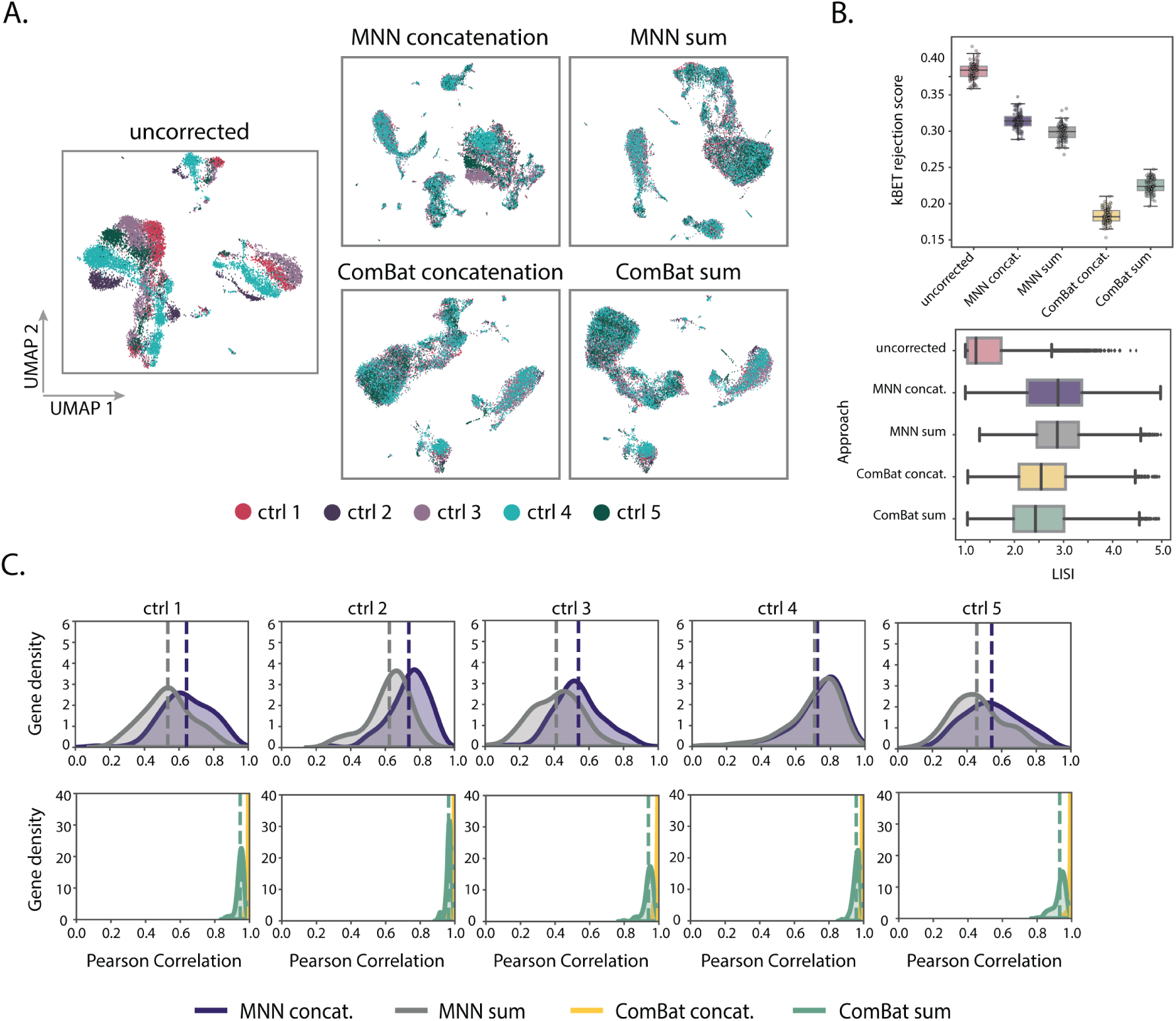
Evaluating batch effect correction for control patient PBMCs in MS case/control dataset. (A) UMAP visualization of control patient PBMCs across batch correction strategies. Spliced and unspliced modalities were combined via concatenation or sum prior to correction with mutual nearest neighbors (MNN) or ComBat. Method performance was measured by batch label mixing metrics kBET and LISI (B), as well as the preservation of the relationship between spliced and unspliced counts (C). Distributions represent the per gene Pearson correlation between cell-cell distances in the phase space (unspliced, spliced) of corrected data and the cell-cell distances in the phase space of each individual control patient. Top panel: Pearson correlation of MNN concatenation or MNN sum to control patients. Bottom panel: Pearson correlation of ComBat concatenation or ComBat sum to control patients. Dashed line represents the mean correlation.

**Supplementary Figure 10:**
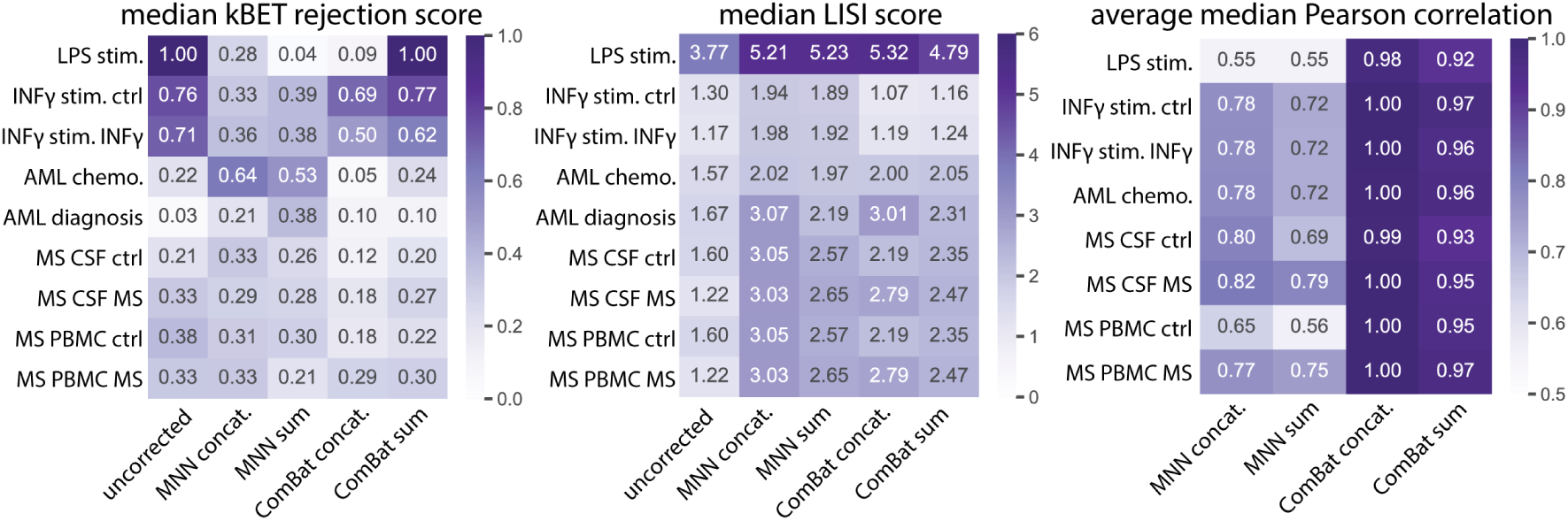
Overall performance of batch correction approaches across perturbation and disease datasets. Batch effect correction performance was assessed according to three metrics, including the median kBET rejection score, median LISI score, and average median Pearson correlation of phase space distances. A correction approach was selected for each dataset if it had the lowest kBET score, highest LISI score, and highest Pearson correlation score.

**Supplementary Figure 11:**
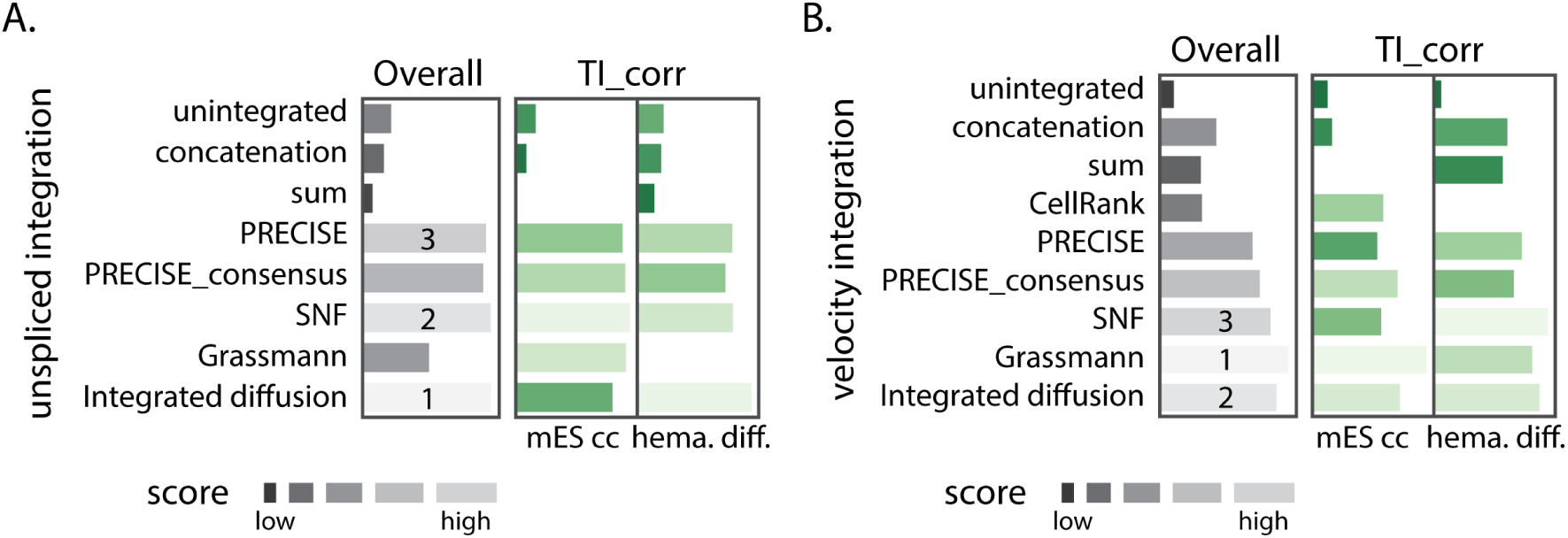
Ranked integration method performance for trajectory inference. Integration methods were ranked by their performance on inferring biological trajectories across mouse embryonic stem cell cycle (mES cc) and mouse hematopoiesis differentiation (hema. diff.) datasets. Individual methods were first ranked according to a trajectory inference correlation (TI_corr_) score, which measures the harmonic mean of cellular positioning correlation and feature importance score correlation to a ground truth reference. The overall performance was then assessed by taking the average of ranked scores across datasets. (A) Overall quality of spliced and unspliced integration performance on inferring biological trajectories. (B) Overall quality of moments of spliced and RNA velocity integration performance on inferring biological trajectories. Here, a higher score is represented by a longer lighter bar. Across both datasets and modalities, intermediate integration approaches outperform unintegrated data on trajectory inference.

**Supplementary Figure 12:**
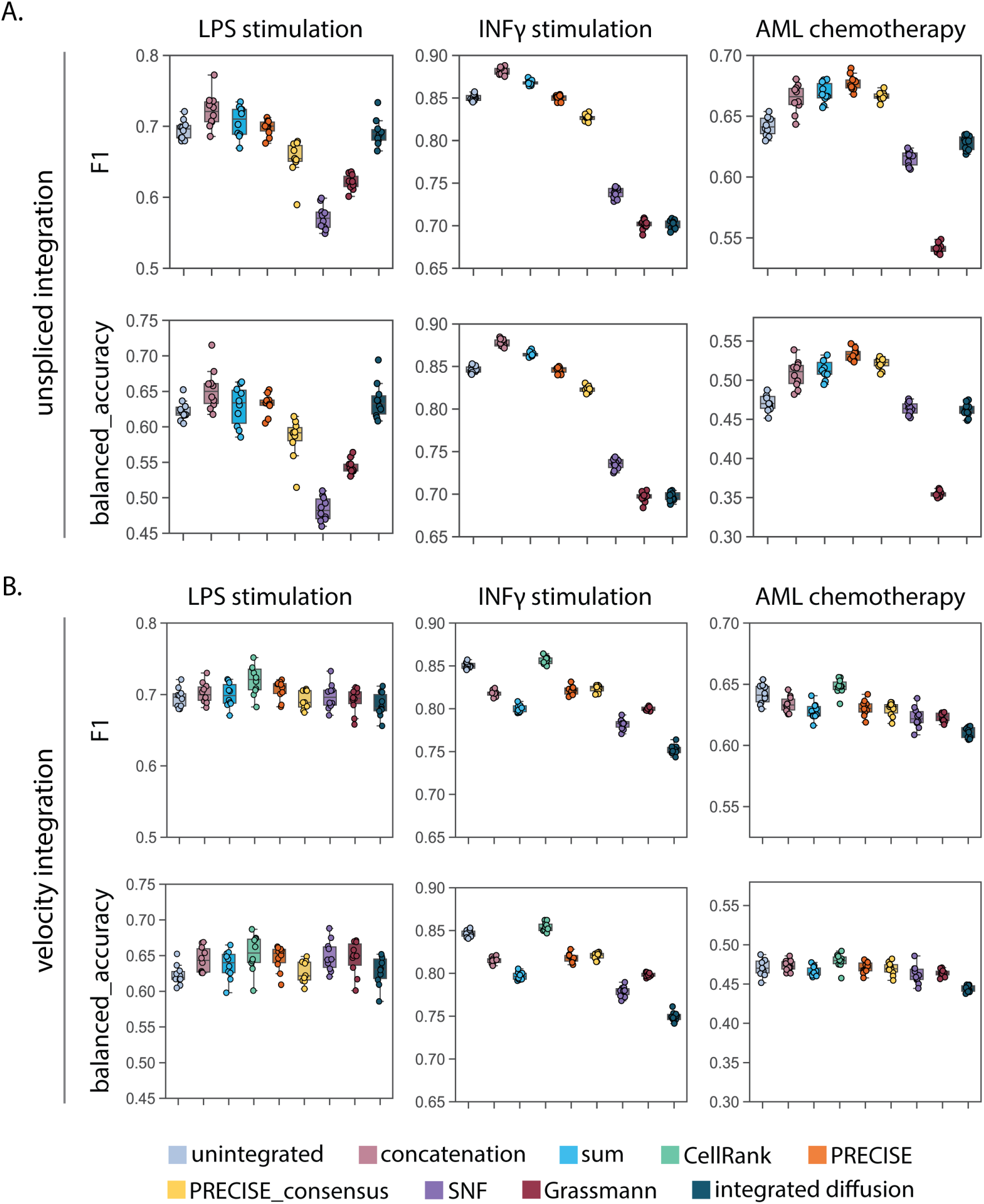
Integration performance on classifying cells according to perturbation condition labels using label propagation. Label propagation was used to classify cells according to treatment condition from (A) spliced and unspliced or (B) moments of spliced and RNA velocity integrated features generated from eight integration approaches. The boxplots represent classification accuracy according to two metrics, top panel: F1 score, bottom panel: balanced accuracy score.

**Supplementary Figure 13:**
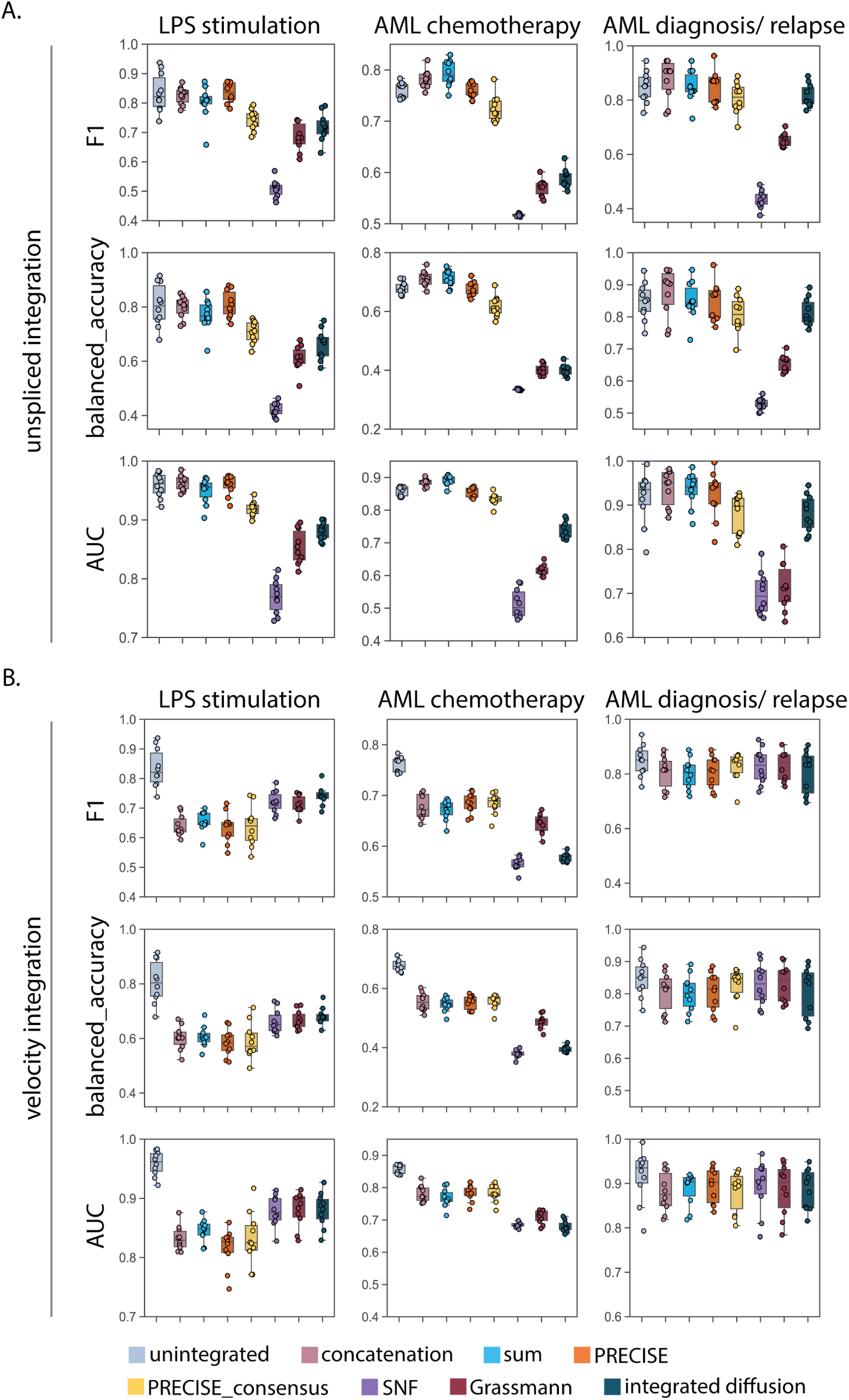
Integration performance on classifying cells according to perturbation condition or disease status using a support vector machine (SVM) classifier. A SVM classifer was used to classify cells according to treatment condition or disease status from (A) spliced and unspliced or (B) moments of spliced and RNA velocity integrated features generated from eight integration approaches. The boxplots represent classification accuracy according to three metrics, including a F1 score, balanced accuracy, and area under the receiver operator curve (AUC).

**Supplementary Figure 14:**
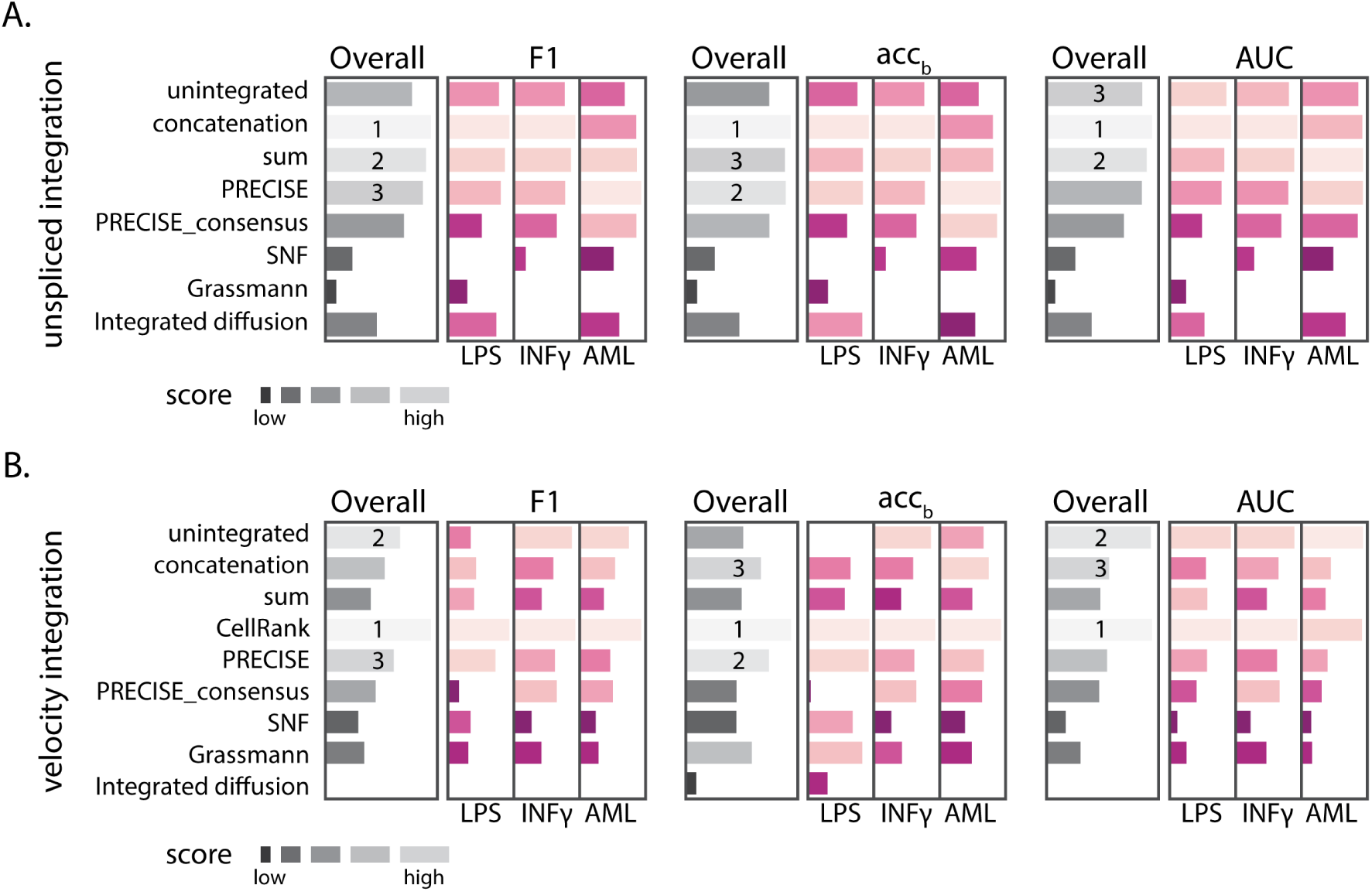
Ranked integration method performance for perturbation classification. Integration methods were ranked according to their performance on classifying cells according to perturbation condition across three datasets, including LPS stimulation of macrophage-like cells (LPS), INF*γ* stimulation of pancreatic islet cells (INF*γ*), and chemotherapy treated cells from a patient with Acute Myeloid Leukemia (AML). Label propagation was used to classify cells according to treatment condition and methods were evaluated by computing three metrics of success: F1 score, balanced accuracy (acc_b_), and area under the receiver operator curve (AUC). The overall performance was then assessed by taking the average of ranked scores across datasets for each metric. (A) Overall quality of spliced and unspliced integration performance on classification of treatment condition. (B) Overall quality of moments of spliced and RNA velocity integration performance on classification of treatment condition. Here, a higher score is represented by a longer lighter bar. Across all three datasets and metrics, spliced and unspliced integration with concatenation and sum outperformed unintegrated data on perturbation classification.

**Supplementary Figure 15:**
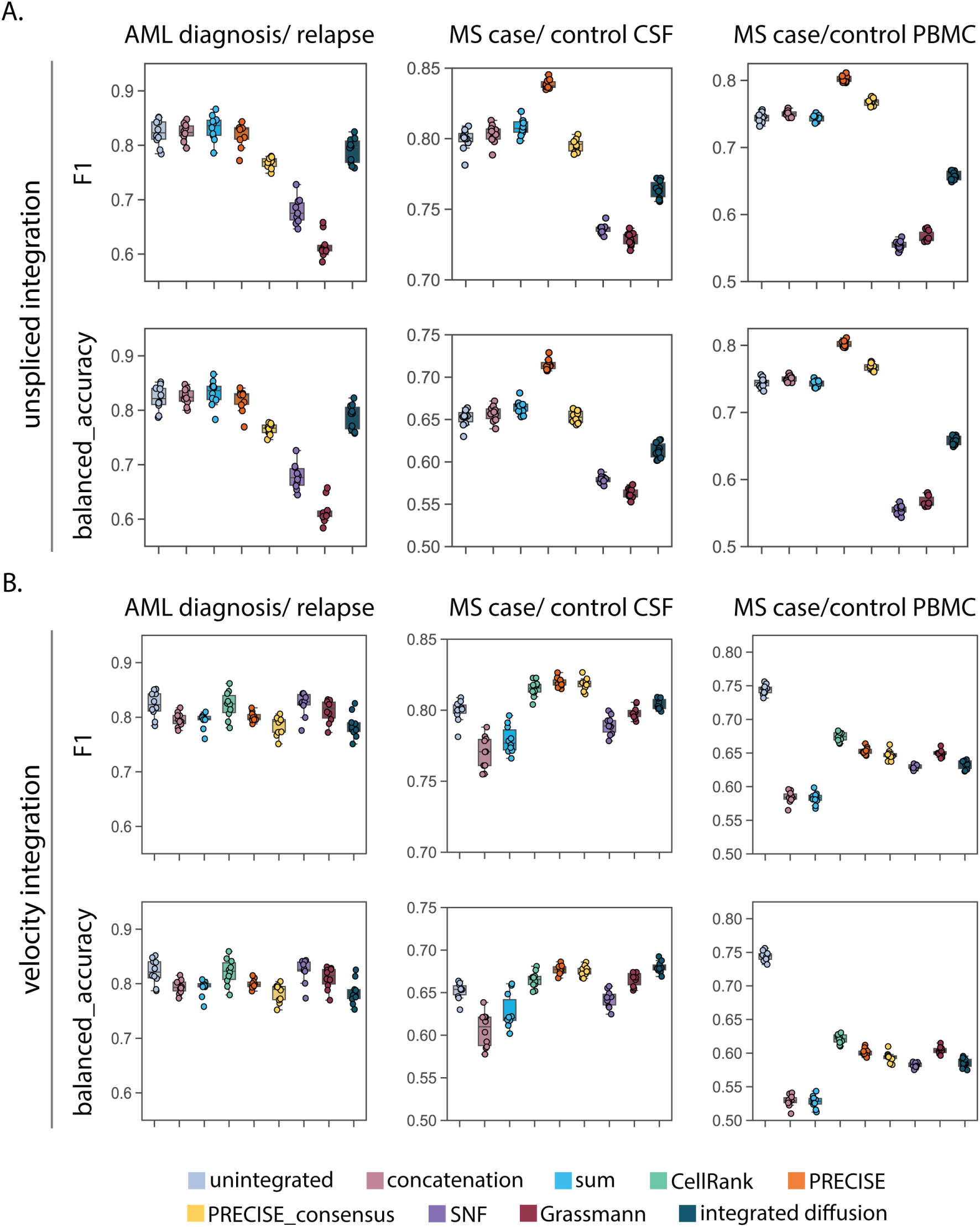
Integration performance on classifying cells according to patient disease status using label propagation. Label propagation was used to classify cells according to patient disease status from (A) spliced and unspliced or (B) moments of spliced and RNA velocity integrated features generated from eight integration approaches. The boxplots represent classification accuracy according to two metrics, top panel: F1 score, bottom panel: balanced accuracy.

**Supplementary Figure 16:**
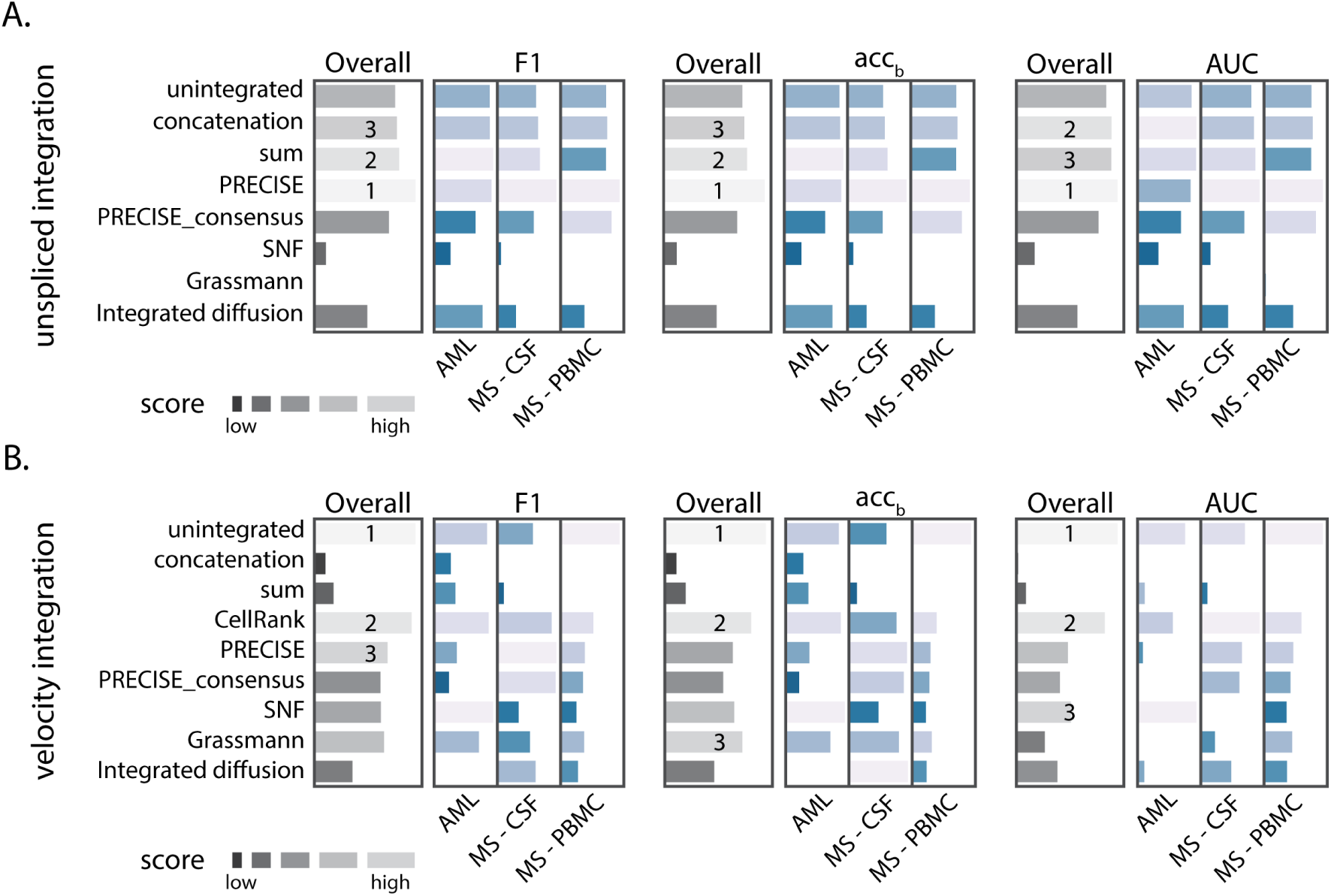
Ranked integration method performance on disease state classification. Integration methods were ranked according to their performance on predicting whether cells were from a healthy or disease patient across three datasets, including an Acute Myeloid Leukemia diagnosis and relapse dataset (AML), a Multiple Sclerosis case/control dataset of cerebral spinal fluid (MS-CSF), and a Multiple Sclerosis case/control dataset of peripheral blood mononuclear cells (MS-PBMC). Label propagation was used to classify cells according to patient disease status and methods were evaluated by computing three metrics of success: F1 score, balanced accuracy (acc_b_), and area under the receiver operator curve (AUC). The overall performance was then assessed by taking the average of ranked scores across datasets for each metric. (A) Overall quality of spliced and unspliced integration performance on classification of cells according to patient disease status. (B) Overall quality of moments of spliced and RNA velocity integration performance on classification of cells according to patient disease status. Here, a higher score is represented by a longer lighter bar. Across all three datasets and metrics, spliced and unspliced integration with PRECISE, concatenation and sum outperformed unintegrated data on disease state prediction.

